# Visualization of photoreceptor outer segment renewal using AAV-delivered Dendra2-tagged rhodopsin

**DOI:** 10.1101/2025.11.30.691349

**Authors:** Lijing Xie, Dianlei Guo, Xiaoyu Zhang, Yingchun Su, Lei Shi, Shujuan Xu, Rong Ju, Tiansen Li, Chunqiao Liu

## Abstract

Visualization of photoreceptor outer segment (OS) renewal dynamics is essential for vision research but has proved difficult in mice due to the small size and dense packing of their photoreceptors. Lack of effective protein “trackers” and the time-consuming generation of transgenic reporter lines add to these challenges. In this study, we evaluated AAV-mediated delivery of photoconvertible Rhodopsin/Dendra2 and Peripherin2/Dendra2 fusion proteins as a means to track OS renewal in mouse photoreceptors. OS renewal was assessed by two approaches: (1) comparing the lengths of native (green) Dendra2 domains at two post-infection time points, and (2) using photoconversion of Dendra2 to distinguish newly synthesized discs from preexisting ones within the same cell. We validated this method in both wild type mice and a *Tmem138-*deficient ciliopathy mutant, in which a reduced OS renewal rate was observed. Taken together, this method offers a rapid genetic tool for “real-time” evaluation of OS renewal dynamics in mice, overcoming limitations posed by the compact and densely organized photoreceptor architecture.

## Introduction

The photoreceptor outer segment (OS) is the initial site of light reception, initiating the image-forming process through a phototransduction cascade. Each rod photoreceptor OS contains approximately 1,000 tightly stacked membranous discs, which are non-contiguous with the surrounding plasma membrane and are densely packed with rhodopsin, the light sensing photopigment. Photoisomerization of 11-*cis* to all-*trans* retinal triggers a conformational change in rhodopsin, initiating the phototransduction cascade.

Approximately 10% of the OS is renewed daily through phagocytosis of shed older discs at the tip by the retinal pigment epithelium (RPE) ^1–4^ and addition of new discs at the base. Renewal requires active transport of newly synthesized proteins and lipids from the biosynthetic inner segment (IS) to the base of the OS. During this process, the plasma membrane at the base of the OS, just distal to the connecting cilium, evaginates to form nascent open discs, which subsequently extend to the full width of the OS and fuse at the edges^5–7^ thereby pinching off from the plasma membrane of the OS. Rhodopsin, the most abundant OS protein, constitutes ∼80% of its total protein mass^8^, providing both structural and functional support for OS structure and phototransduction. Rhodopsin is confined to each mature disc, as it cannot diffuse between physically separated disc membranes.

Therefore, the domain occupied by newly synthesized rhodopsin largely represents the nascent discs at the base.

Rhodopsin transport can be broadly divided into two steps: 1) During post-Golgi trafficking rhodopsin-carrying vesicles are transported to the photoreceptor ciliary base, a process coordinated by GTPases, their effectors, intraflagellar transport (IFT) complexes, molecular motors^9–11^, and rhodopsin C-terminal targeting motifs (e.g., VXPX) ^9, 12–15^; 2) Delivery across the connecting cilium of rhodopsin to the OS discs requires a switching of molecular motors and IFT complexes at the ciliary base^16^. Mutations in IFT B subunits or associated kinesin motors often lead to rhodopsin mislocalization^17, 18^.

The photoreceptor ciliary gate plays a critical role in OS rhodopsin transport. The OS is a specialized sensory cilium that relies on the connecting cilium, analogous to the transition zone of primary cilia, and associated protein complexes to function as a diffusion barrier to selectively transport proteins to the OS^9, 19, 20^. Disruption of key ciliary gate components, including the NPHP, MKS, JBTS, and BBS protein modules, causes photoreceptor degeneration characterized by rhodopsin mislocalization^21–24^.

Given its sheer abundance, rhodopsin also serves a structural role in disc membrane homeostasis^25^. By the same token, mislocalization of rhodopsin is bound to have a deleterious impact in the organelles or membrane domains where it is trapped. Despite the importance of correct rhodopsin localization in photoreceptor structure and function and in disease pathophysiology^26–29^, determining rhodopsin OS transport defects is often infeasible due to the lack of means to monitor this dynamic process as it progresses. Secondary effects from failed OS development or structural disruptions further confound data interpretations. In addition, a transport machinery usually carries multiple cargos, and defective ciliary gates can also allow IS-localized proteins to leak into the OS, causing rhodopsin mislocalization and photoreceptor degeneration^30^.

The photoconvertible fluorescent protein-tagging technique is a valuable tool for distinguishing newly synthesized proteins from pre-existing ones. Dendra2 is a newer variant of photoconvertible fluorescent Dendra, originally derived from *Dendronephthya sp.* with a single amino acid substitution (A224V) and higher photoconversion efficiency, photostability and brightness. Unlike many other fluorescent proteins, Dendra2 is monomeric and relatively small (26kd), thus suitable for directly tagging and tracing a specific protein. It can be efficiently photoactivated with ultraviolet illumination, leading to irreversible cleavage and reformation of covalent bonds with ensuing spectral shift from green to red^31, 32 33^ (Suppl. Fig. 1). This feature is particularly useful for studying the renewal of spatially and temporally isolated structures, such as OS discs. Previously, a Dentra2 tagging-based method has been employed successfully to study protein transport in *Xenopus* photoreceptors^15, 34–38^. It has also been adapted to investigate various cellular events in mice^39–41^, but not in the photoreceptor cells. Since mouse photoreceptors are five times smaller in diameter compared to *Xenopus* ^42^, applying methodologies developed in Xenopus to the mammalian retina are expected to meet technical challenges and will require developing new methods uniquely suited to the much smaller mammalian photoreceptors.

To develop useful tools for studying protein trafficking and OS renewal, we utilized photoreceptor-targeting AAV vectors to express photoconvertible fluorescence-tagged rhodopsin and peripherin 2 (formerly known as RDS) in the mouse retina^43^. For validation, we included both wild type and a ciliopathy mutant line with disruption of *Tmem138*, previously shown to be important for OS morphogenesis^24^. Our data show that the rate of OS renewal can be quantified readily with the method, and differences between wild type and the trafficking defective mutant can be distinguished with this approach. This study demonstrated proof of principle and provides a novel and valuable tool for tracking and differentiating OS renewal rates in both wild type and mutant mice with defects in OS morphogenesis and renewal.

## Results

### Rhodopsin/Dendra2 photoconversion *in vitro*

We first validated the photoconversion capacity of the Rhodopsin/Dendra2 fusion protein by constructing a lentiviral vector encoding the *Rhodopsin/Dendra2* gene under the control of the CMV promoter and transducing it into 293T cells (Fig. 1A). Forty-eight hours post-transduction, green fluorescence from Rhodopsin/Dendra2 expression was readily detected by confocal microscopy (Fig. 1B, left column). Subsequent scanning with a 488 nm laser line for 10 min did not produce detectable red spectral shift/photoconversion (Fig. 1C), indicating spectral stability of Dendra2 under standard green-channel imaging conditions, thus preventing ambiguity that may arise when simultaneously imaging green and red-shifted forms of Dendra2.

**Figure 1.**
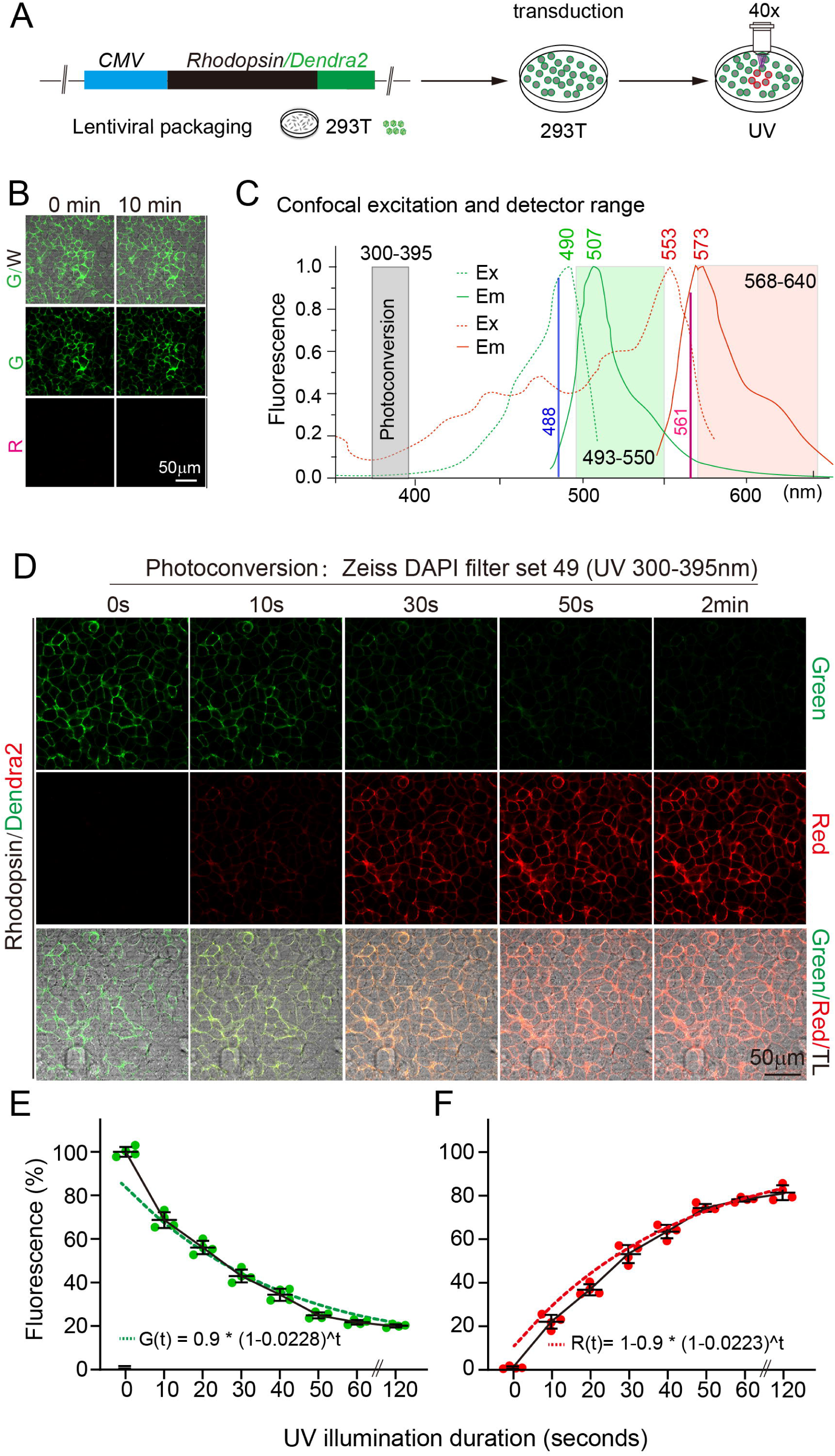
Rhodopsin/Dendra2 photoconversion in cultured HEK293T. (**A**) Schematic overview of the experimental workflow: Lentiviral Rhodopsin/Dendra2 construction, viral packaging, transduction of 293T cells, and photoconversion using UV light. UV spectra were generated using a mercury arc lamp (X-Cite 120Q, EXCELITAS) equipped with Zeiss LSM880 confocal microscopy bandpass filtered for 300–395 nm using Zeiss DAPI filter set 49 (excitation: G365; beam splitter: FT395; emission: BP445/50). (**B**) Expression of Rhodopsin/Dendra2 in 293T cells. Note that 10 minutes of 488nm laser exposure did not induce photoconversion of Dendra2. (**C**) Photoconversion and imaging settings according to Dendra2 chromophore properties. The gray box represents UV spectra used for photoconversion. Dashed lines and solid lines indicate excitation (Ex) and emission (Em) spectra respective to Dendra2 before (green) and after (red) photoconversion (remade from Chudakov et al., 2007, reference 42).Green fluorescence was excited using a 488 nm confocal laser line and collected with a photon detector banded for 493–550 nm (greenish box), while the shifted red spectra was excited by 561nm laser and collected with a photon detector set for 568–640 nm range (pinkish). (**D**) UV irradiation conducted at 164 lux (39 μw /cm^2^) through a 40x lens. First row: green fluorescence signal from the residual unconverted Dendra2 fluorescence along the indicated time axis with UV radiation; Second row: increased converted red fluorescence signal; Third row: merged red, green, and transparent light (TL) channels demonstrating a negative correlation of the red and green fluorescence given the irradiated time. (**E, F**) Plotted ImageJ measurements (n= 4 images) of green and red relative intensities along the time axis of radiation at 10-second intervals. Dashed lines represent exponential fits to the data. The green fluorescence decay was fit with the function: *G(t)* = 0.9·(1 – 0.0228)□, while the red fluorescence increase followed: *R(t)* = 1 – 0.9·(1 – 0.0223)□.

We next determined the time-course of UV photoconversion by exposing the cells to UV light in the wavelength range of 300–395nm (with peak at 365 nm) following previous studies^44^. After photoconversion, green fluorescence was excited using a 488 nm confocal laser line and collected with a photon detector banded for 493–550 nm, while the red-shifted spectra were excited by 561 nm laser and collected with a photon detector set for 568–640 nm range (Fig. 1C). Progressive decline of green fluorescence with concomitant increase of red fluorescence was observed over time (Fig. 1D). Notably, about 20% of green fluorescence remained unconverted even as red fluorescence intensity peaked (Fig. 1D, last panel of the top row; Fig. 1E, F). The residual green fluorescence was not due to reversion, as the red shift of Dendra2 is irreversible, involving covalent bond breakage, reformation and loss of functional groups (Suppl. Fig. 1A). Nor could it be explained by bleed-through from other channels given the set imaging conditions. This observation was, however, consistent with a previous study that found photoconversion of Dendra2 to be incomplete^32^.

We next performed a quantitative analysis of relative fluorescence intensities. Average fluorescence intensities from four imaging fields were measured by NIH ImageJ at eight time points with UV illumination applied at 10-second increments (Table 1, Fig. 1E, F). Measured green fluorescence was normalized to the initial green intensity, and the red fluorescence, regardless of its absolute value, was normalized to 0.8 of the initial green, assuming a 20% residual green signal and all photoconverted Dendra2 contributing to the red emission (Fig. 1D). While this simplified approach ignores possible differences in extinction coefficients and quantum yields between the red and green forms of Dendra2, it nevertheless demonstrates that both the decay of green fluorescence and the increase in red fluorescence closely followed exponential fits, with comparable calculated rate constants (0.0228 vs. 0.0223 s⁻^1^, respectively) (Fig. 1E, F), supporting a time-dependent photoconversion process.

**Table 1.**
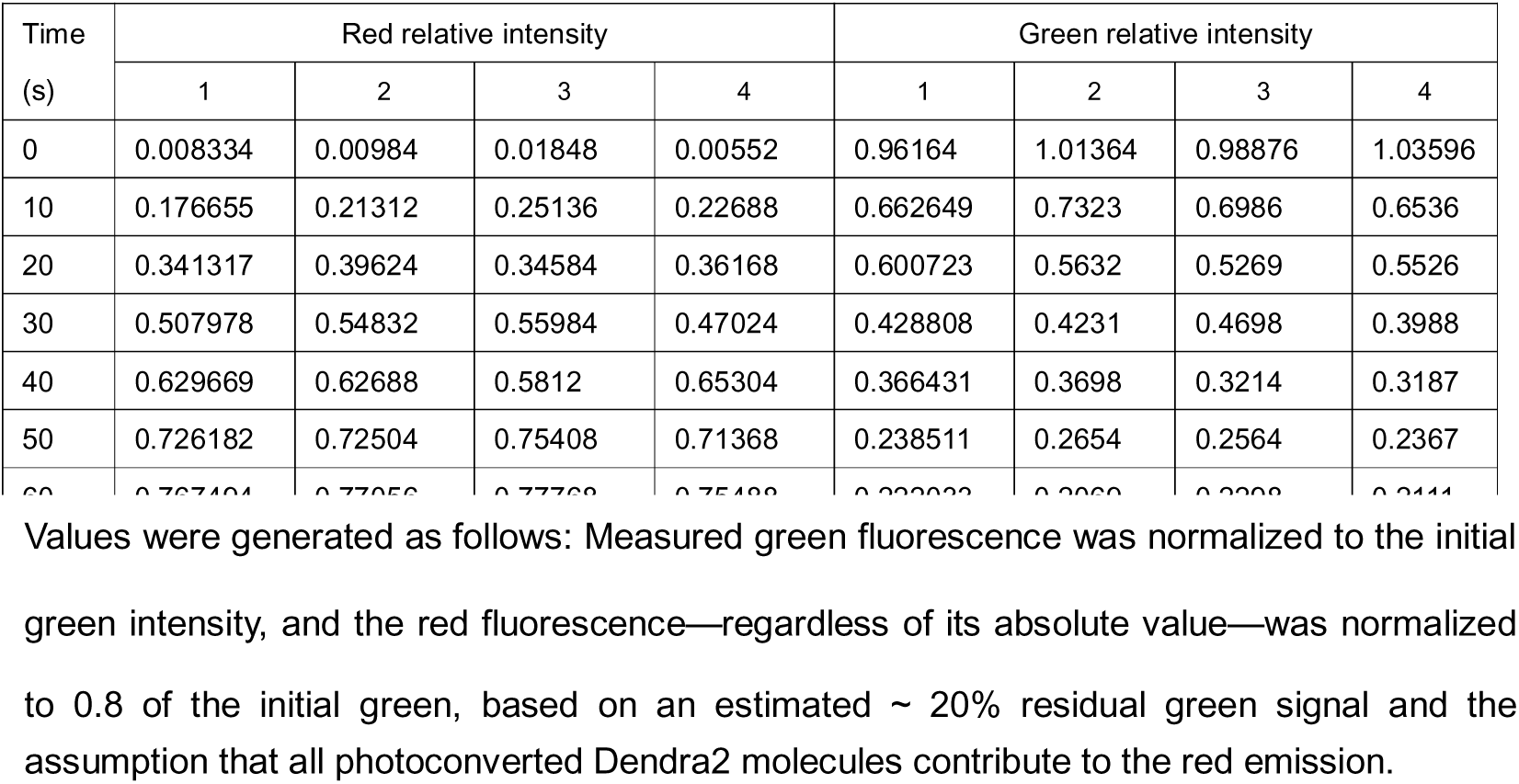
Measured fluorescence intensities from four imaging fields of unconverted green and converted red Dendra2 forms.

### Optimization of photoconversion Rhodopsin/Dendra2 *in vivo*

Having validated it *in vitro*, we went on to examine the photoconversion of Rhodopsin/Dendra2 in the mouse retina by adeno-associated viral (AAV) vector-mediated gene delivery. We constructed an AAV vector in which expression of Rhodopsin/Dendra2 was driven by a photoreceptor-specific promoter (Fig. 2A) and injected the vector into the subretinal space of mice (Fig. 2B–D). Ten days post-injection, mice were anesthetized, pupils dilated and placed on the stage of a Zeiss ApoTome upright epifluorescence microscope for UV irradiation of their fundi at 120 lux / 31 µW/cm² for 30 minutes using a 4× objective lens (Fig. 2E). Mice were positioned such that the illuminated retinal quadrants were primarily nasal. Fundus images were acquired using Zeiss green and red filter sets to minimize channel bleed-through (Fig. 2F, G). Photoconverted red fluorescence was clearly visible alongside unconverted green fluorescence (Fig. 2H–J). Subsequent analysis of flat-mount retinas found photoconverted red fluorescence in the UV illuminated areas (Fig. 2K), while the area without UV exposure showed only green fluorescence (Fig. 2L).

**Figure 2.**
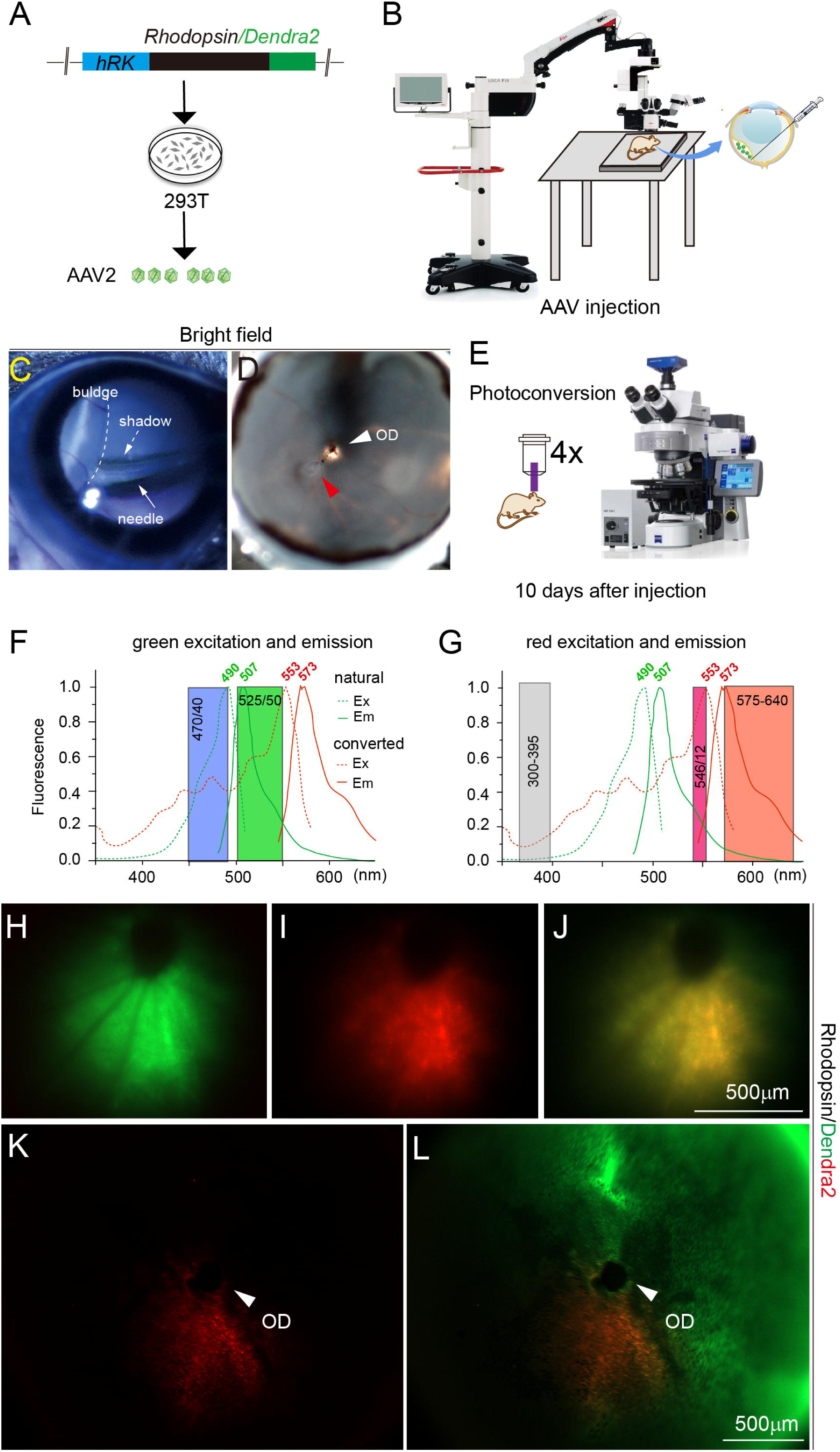
Rhodopsin/Dendra2 photoconversion observed by fundus fluorescence. (**A**) Construction and production of AAV2 viral particles carrying *Rhodopsin/Dendra2* driven by a rod-specific human rhodopsin kinase (hRK) promoter in 293T cells. (**B**) Schematic illustration of purified viral particles injection into the subretinal space of the mouse retina under an ophthalmic microscope. (**C**) A fundus picture showing the injection needle (arrow) and a bleb (dashed lines) caused by temporary retinal detachment in an injected eye. The dashed arrow points to the shadow of the injection needle. (**D**) Fundus of a 4% PFA-fixed eye cup after removal of the lens and cornea 10 days after viral infection. The red arrowhead points to the injection site, and the white arrowhead points to the optic disc. (**E**) Radiating and imaging the mouse retina infected by AAV viral particles expressing *Rhodopsin/Dendra2* 10 days after AAV infection through 4x objective. (**F, G**) Dashed lines and solid lines indicate excitation (Ex) and emission (Em) spectra respective to Dendra2 before (green) and after (red) photoconversion. Boxes in (**F**) indicate filter sets for excitation (470/40) and emission (525/50) of Dendra2 green form. Boxes in (**G**) indicate filter sets for excitation (546/12) and emission (575-640) of photoconverted Dendra2. Gray box: Isolated UV spectra of 300-395nm using DAPI filter set 49: G365; FT395; BP445/50 for photoconversion. (**H**) Fundus picture of unconverted green fluorescence. (**I**) Fundus was UV radiated for 30min viewed through a red emission filter. (**J**) A merged fundus image from residual green and converted red fluorescence. (**K**) A dissected flat mount retina fixed with 4% PFA showing photoconverted red fluorescence. (**L**) A merged image from residual green and converted red fluorescence of the same retina as in (**K**). Arrowheads point to the optic discs. Note the green fluorescence in area without UV radiation.

We next sought to optimize the conditions for *in vivo* photoconversion by varying UV intensity levels and exposure durations. Using the 5-step intensity generator (Materials and Methods), we performed illumination through a 4× objective lens at 40 lux / 5 µW/cm² (Level I), 66 lux / 18 µW/cm² (Level II), 120 lux / 31 µW/cm² (Level III), and 390 lux / 96 µW/cm² (Level IV). At each power level, we tested three exposure durations at 5, 10, and 15 minutes. Images were acquired using the same filter sets as described in Figure 2. Fundus imaging showed increased red fluorescence with higher illumination powers or longer exposure times (Fig. 3A-H). The intensity and time dependence of photoconversion were further quantified from four imaging fields of four individual eyes (Fig. 3I-L).

**Figure 3.**
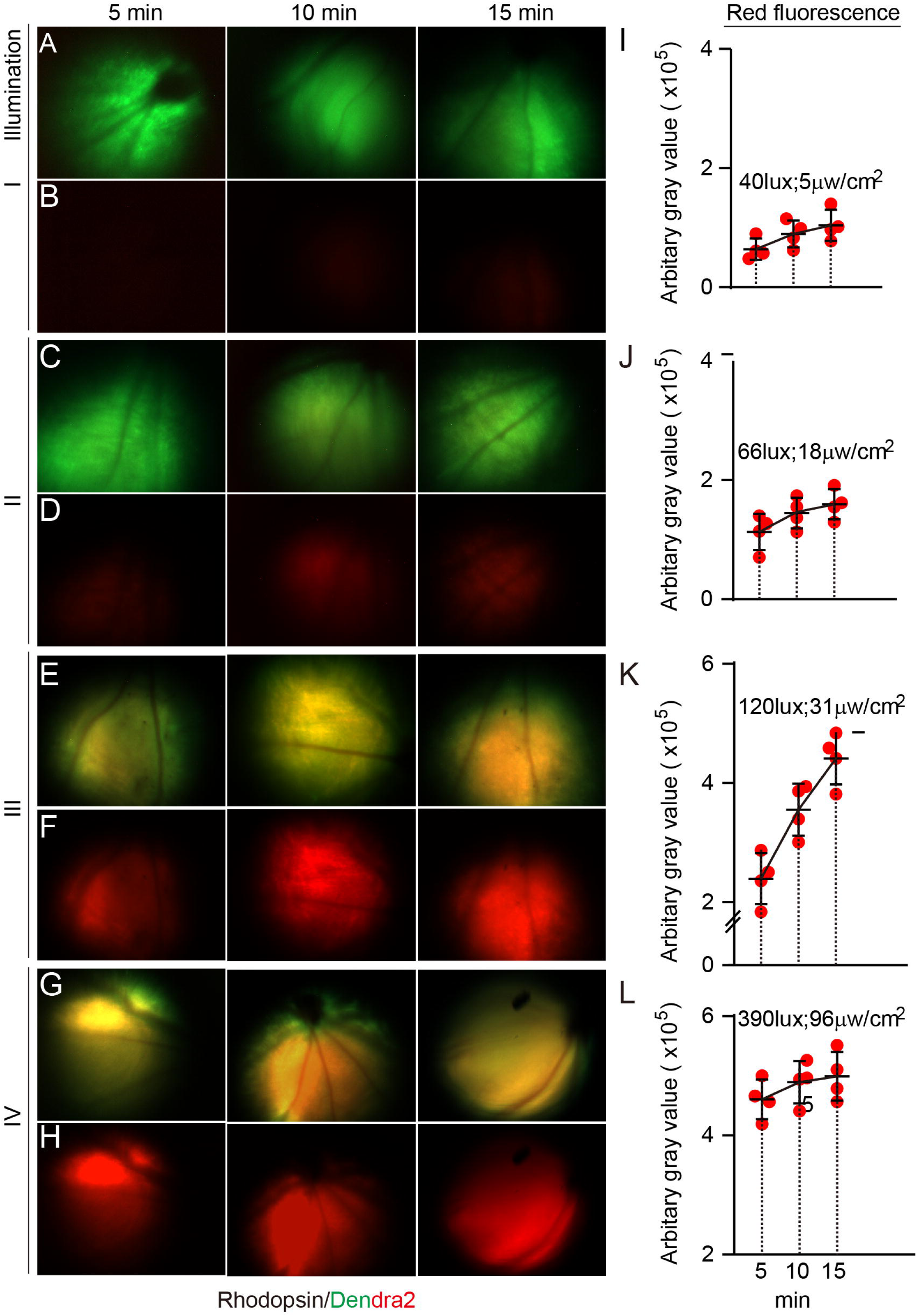
Optimization photoconversion illuminance by fundus fluorescence imaging. Fundus images were acquired following exposure to blue light at varying intensities (Levels I–IV) for 5, 10, and 15 minutes. Level I, 40 lux /5 μw /cm^2^; Level II, 66 lux /18 μw/cm^2^; Level III,120 lux /31 μw/cm^2^; Level IV, 390 lux / 96 μw/cm^2^. Image collection filter sets were described in the previous figure. (**A, B**) Fundus images under Level I illuminance at three time points. (**A**) Merged red and green fluorescence channels. (**B**) Red fluorescence channel only. (**C**, **D**) Images from Level II illuminance, displayed as in panels (**A**, **B**). (**E**, **F**) Images from Level III illuminance, arranged as in panels (**A**, **B**). (**G, H**) Images from Level IV illuminance, arranged as in panels (**A**, **B**). (**I-L**) Quantification of fluorescence intensities from four imaging fields (4 eyes). Fluorescence values are measured using NIH Image J and are presented in arbitrary units (AU).

Photoconverted fluorescent proteins were further analyzed at higher resolution on retinal sections by confocal microscopy following Level II and III UV illumination, along with the endogenous rhodopsin from untransduced photoreceptors detected by the ID4 antibody (Fig. 4A-F). At higher magnification, Rhodopsin/Dendra2 faithfully recapitulated endogenous rhodopsin localization (Fig. 4G, G’, H, H’, H’’). Similar photoconversion was observed in a photoreceptor-defective *Tmem138* mutant retinas (Fig. 4I, I’, J, J’, J’’), with inner segment mislocalization of Rhodopsin/Dendra2 ^24^. We also assessed the extent of light damage of Level III and Level IV illumination over time and found only minimal retinal damage reflected by GFAP-stained glial activation and DAPI-stained retinal structure, 2 days after illumination (Suppl. Fig. 2). Overall, Rhodopsin/Dendra2 underwent readily detectable levels of photoconversion *in vivo*, and Level III UV illumination for a brief duration of 5 minutes appeared to be optimal, taking into account photoconversion efficiency, procedural practicality and minimal retinal light damage.

**Figure 4.**
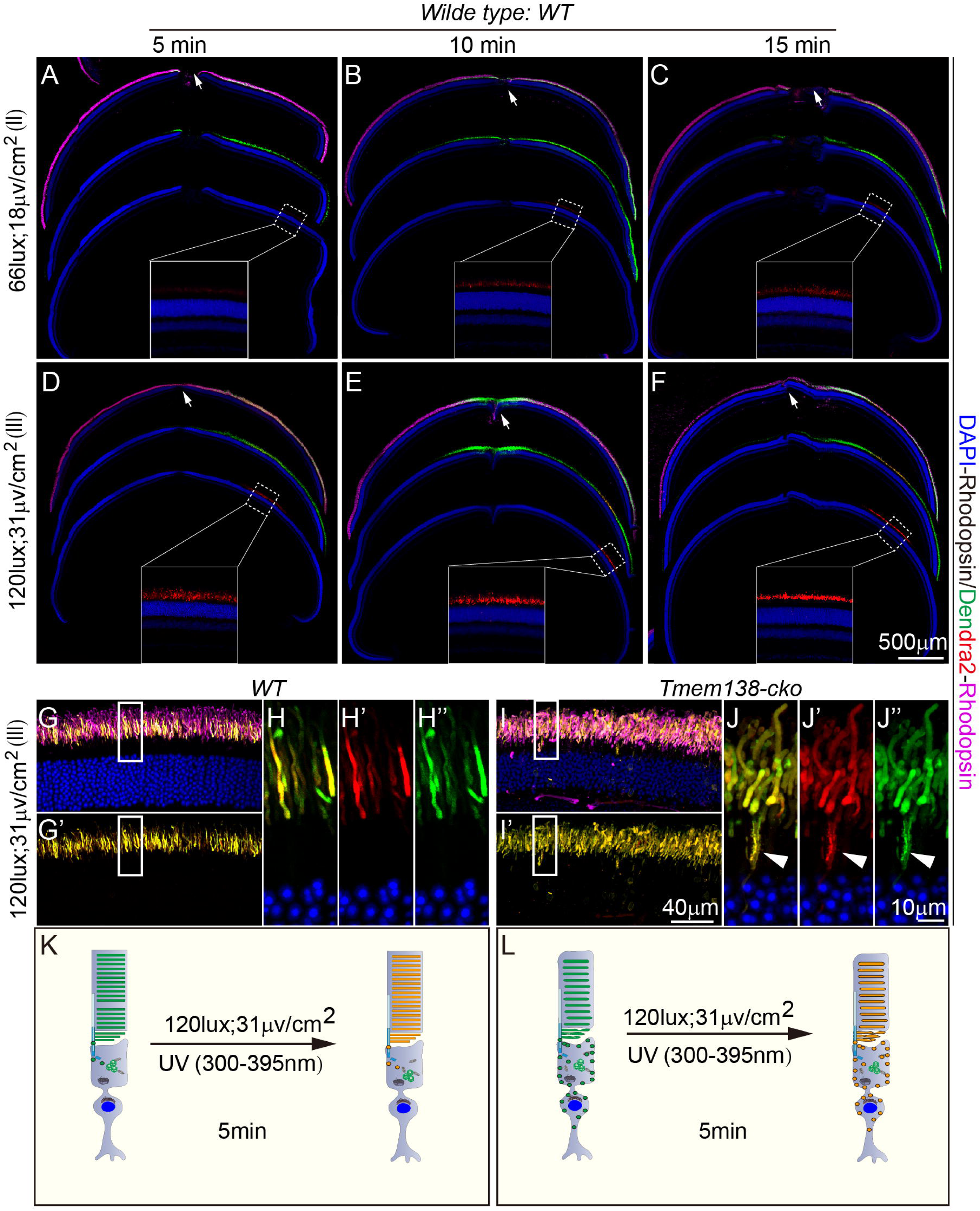
Photoconversion of Rhodopsin/Dendra2 observed in retinal sections at varying levels of illuminance. (**A-F**) Confocal images of retinal sections following UV irradiation at two intensities: 66 lux / 18 μw/cm^2^ (Level III) and 120 lux / 31 μw/cm^2^ (Level IV) for 5, 10, and 15 minutes. Arrows point to the viral injection sites. Mangena represents ID4-stained endogenous rhodopsin. Green and red are unconverted and converted Rhodopsin/Dendra2, respectively. Insets are magnified views of boxed areas showing red fluorescence in converted cells. (**G**) Retinal section from a wild type eye showing merged channels of endogenous rhodopsin (magenta), Rhodopsin/Dendra2 (green), and converted red fluorescence. (**G’**) Merged green and red channels corresponding to (**G**). (**H-H”**) Higher magnification of the boxed region in (**G’**), showing the merged green and red channels (**H**), the red channel alone (**H’**), and the green channel alone (**H**”). (**I, I’; J–J”**) Retinal sections from *Tmem138*-cko mutant mice showing defective outer segments and disrupted rhodopsin transport. Panel arrangement follows that of (**G**), (**G’**), and (**H–H”**): (**I**) merged channels (green, red, and magenta); (**I’**) merged green and red channels; (**J–J**”) magnified boxed area from (**I’**) showing merged (**J**), red (**J’**), and green (**J”**) fluorescence channels. Arrowheads point to the IS mislocalization. **(K, L)** Schematic illustrations of Rhodopsin/Dendra2 photoconversion in wild type **(K**) and *Tmem138*-cko mutant (**L**) photoreceptors.

### Photoreceptor OS renewal indicated by Rhodopsin/Dendra2 in wild type and a trafficking-deficient mouse line

We then set out to evaluate whether Rhodopsin/Dendra2 could serve as a reporter for tracking outer segment (OS) renewal. We first examined Rhodopsin/Dendra2 domain lengths on Day 4 (D4), D6, and D10 following AAV transduction (Fig. 5A) in wild type mouse retinas, with endogenous rhodopsin visualized using ID4 antibody. Elongation of Rhodopsin/Dendra2 domain was observed between D4 and D6 (Fig. 5B, B’, B’’ compared to 5C, C’, C’’), and between D6 and D10 (Fig. 5C, C’, C’’ compared to 5D, D’, D’’).

**Figure 5.**
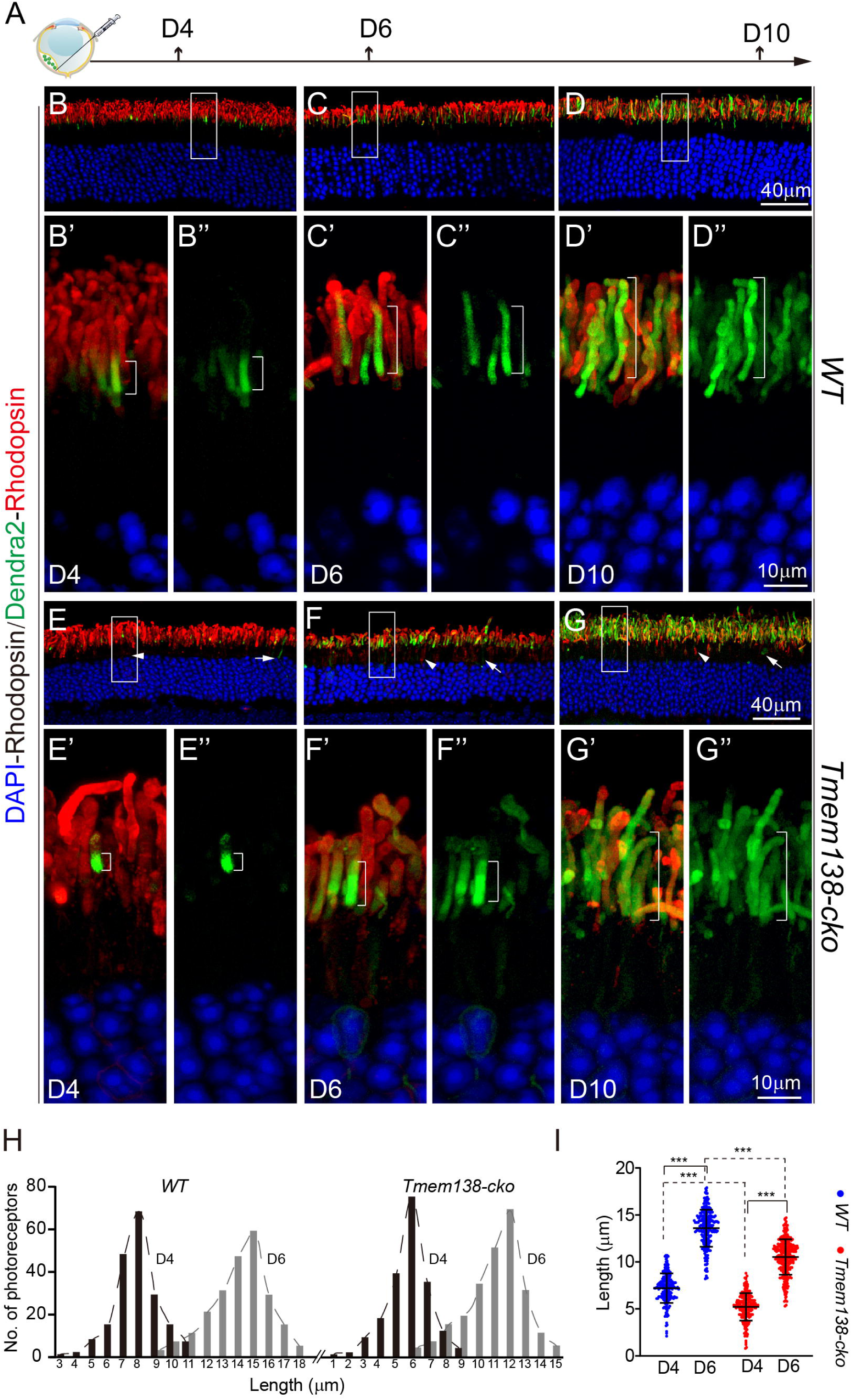
Outer segment renewal revealed by the unconverted Rhodopsin/Dendra2. (**A**) Experimental scheme for detection of Rhodopsin/Dendra2 domain length at different time points after AAV-*Rhodopsin/Dendra2* infection. Samples were collected on day 4 (D4), D6, and D10 post-viral infection. (**B**) Wild type retinal sections of D4 post injection. Shown is a merged image of endogenous rhodopsin (red), detected by the ID4 antibody, and exogenous Rhodopsin/Dendra2 (green) introduced by an AAV viral vector. Nuclei are stained with DAPI (blue). (**B**’, **B**’’) Higher magnification of boxed area in (**B**), displayed as a merged image (**B**’) and the green channel alone (**B**’’) for Dendra2. Brackets indicate the Dendra2 domain within the OS. (**C**, **C**’, **C**’’) Wild type retinal sections at D6 post-injection, shown with the same channel arrangement as in (**B**’, **B**’’). (**D**, **D**’, **D**’’) Wild type retinal sections at D10 post-injection. (**E**) *Tmem138*-*cko* retinal sections at D4 post-injection. The arrowhead points to the mislocalization of rhodopsin (red) in the mutant inner segments; the arrow points to the inner segment Rhodopsin/Dendra2 (green). (**E**’, **E**’’) Higher magnification of the boxed region in (**E**), shown as a merged image (**E**’) and a green channel alone (**E**’’) for Dendra2. Brackets mark the Dendra2 domains within the OS. (**F**, **F**’, **F**’’) *Tmem138*-*cko* retinal sections at D6 post-injection. (**G**, **G**’, **G**’’) *Tmem138*-*cko* retinal sections at D10 post-injection. Arrowheads and arrows in (**E-G**) denote mislocalized endogenous rhodopsin (red) and Rhodopsin/Dendra2 (green), respectively, as in (**E**). (**H**) Distribution Rhodopsin/Dendra2 domain lengths of wild type and *Tmem138*-*cko* photoreceptors at D4 (dark gray boxes) and D6 (light gray boxes) post viral infection. Each box represents 1μm bin width. (**I**) Student’s t-test was used to evaluate the significance of differences between wild type and mutant groups. ***, p<0.001.

We next performed the same experiments on *Tmem138* mutant mice. Our previous study showed that the ciliary membrane protein Tmem138 was required for OS morphogenesis^24^. We postulated that the *Tmem138* mutant would alter OS disc renewal dynamics through affecting rhodopsin transport since Tmem138 likely interacts with rhodopsin^24^, thereby serving as a useful tool to validate this method. Indeed, the newly renewed Rhodopsin/Dendra2 domains were noticeably shorter in *Tmem138* mutant photoreceptors compared to controls (Fig. 5E, E’, E’’ compared to 5B, B’, B’’; Fig. 5F, F’, F’’ compared to 5C, C’, C’’). Nonetheless, both wild type and mutant photoreceptors completed one cycle of OS renewal by D10, as indicated by full-length OS filled by Rhodopsin/Dendra2 fluorescence (Fig. 5D, D’, D’’; Fig. 5G, G’, G’’). The comparable renewal cycle time of mutant OS and wild type mice might result from the shortened OS offsetting the slower renewal rate in the mutant.

We next quantified OS renewal in photoreceptors with clearly identifiable fluorescence domains at D4 and D6 (Suppl. Fig. 3A-H, Table 2 and Suppl. Table 1). The distributions of domain lengths, binned at 1μm intervals, roughly followed bell-shape curves—likely reflecting the asynchronous expression onset—and showed temporal separation in both wild type and *Tmem138* mutant photoreceptors (Fig. 5H). The average fluorescent domain lengths between D4 and D6 significantly differed between the wild type and mutant (Fig. 5J), and the calculated growth rates were 3.19 and 2.65 μm/day for the wild type and *Tmem138* mutant photoreceptors, respectively (Table 2). The slower growth rate of Rhodopsin/Dendra2 domain in mutant photoreceptors was consistent with the reported trafficking defects in the *Tmem138* mutant and the previous observed interaction between Tmem138 and Rhodopsin.

**Table 2.** Distribution of photoreceptors with differential Rhodopsin/Dendra2 domain lengths at D4 and D6 postinfection without photoconversion.

| No. PR<br>Length (μm) | WT_D4 | WT_D6 | *Tmem138-cko*_D4 | *Tmem138-cko*_D6 |
| --- | --- | --- | --- | --- |
| 0-1 | 0 | 0 | 2 | 0 |
| 1-2 | 0 | 0 | 3 | 0 |
| 2-3 | 2 | 0 | 10 | 0 |
| 3-4 | 3 | 0 | 19 | 0 |
| 4-5 | 9 | 0 | 40 | 0 |
| 5-6 | 16 | 0 | 76 | 5 |
| 6-7 | 49 | 0 | 36 | 8 |
| 7-8 | 69 | 0 | 13 | 16 |
| 8-9 | 30 | 4 | 5 | 20 |
| 9-10 | 16 | 8 | 0 | 35 |
| 10-11 | 8 | 12 | 0 | 52 |
| 11-12 | 0 | 22 | 0 | 70 |
| 12-13 | 0 | 32 | 0 | 32 |
| 13-14 | 0 | 48 | 0 | 12 |
| 14-15 | 0 | 60 | 0 | 6 |
| 15-16 | 0 | 30 | 0 | 0 |
| 16-17 | 0 | 16 | 0 | 0 |
| 17-18 | 0 | 6 | 0 | 0 |
| Mean | 7.215 μm | 13.59 μm | 5.221 μm | 10.52μm |
| Sd | 1.541 | 1.969 | 1.461 | 1.884 |
| Renewal rate | 3.19μm/day |  | 2.65μm/day |  |

The left column number ranges are binned at 1μm intervals. No. PR, number of photoreceptors ; Sd: Standard deviation

### Photoreceptor outer segment renewal indicated by Rhodopsin/Dendra2 photoconversion

Photo convertibility of Rhodopsin/Dendra2 is a key element that helps ensure a more accurate measurement of OS renewal. A limitation of the measurements based on native green emission of Dendra2 is that OS renewal was inferred from the differences in lengths of the fluorescence domains at two time points after viral infection. This approach was subject to inherent variability in the time to onset and expression levels of Rhodopsin/Dendra2. Photoconverting Rhodopsin/Dendra2 to its red spectra when expression has plateaued allows newly emerging discs (unconverted, green) to be distinguished from pre-existing ones (converted, red) thereby ‘resetting’ the renewal timeline.

Using this strategy, we performed photoconversion at 10 days post -AAV transduction when the OS had been filled by Rhodopsin/Dendra2 along its entire length (Fig. 6A). We applied the previously optimized illumination conditions to the retina, i.e., Level III UV power for 5 minutes (see Fig. 4). Due to the presence of unconverted Rhodopsin/Dendra2, OS domains exhibiting both green and red fluorescence were considered old discs, whereas regions displaying purely green fluorescence were newly formed discs (Fig. 6B-E, B’-E’). OS renewal was barely detectable 6 hours after photoconversion (insets of Fig. 6B, B’, C, C’) but by 48 hours post-photoconversion, pure green fluorescence-labeled domains representing new discs were clearly visible at the base of the OS in both wild type and *Tmem138* mutant photoreceptors (insets of Fig. 6D, D’, E, E’, brackets). Quantification of the green fluorescent domains after photoconversion revealed renewal rates of 2.9µm/day (sd=0.2565) in wild type and 2.4µm/day (sd=0.3698) in *Tmem138* mutant mice (Table 3, Suppl. Table 1, Fig. 6F, G). As expected, measurements of OS renewal based on photoconversion exhibited lower variability compared to measurements obtained without photoconversion (Table 2, 3).

**Figure 6.**
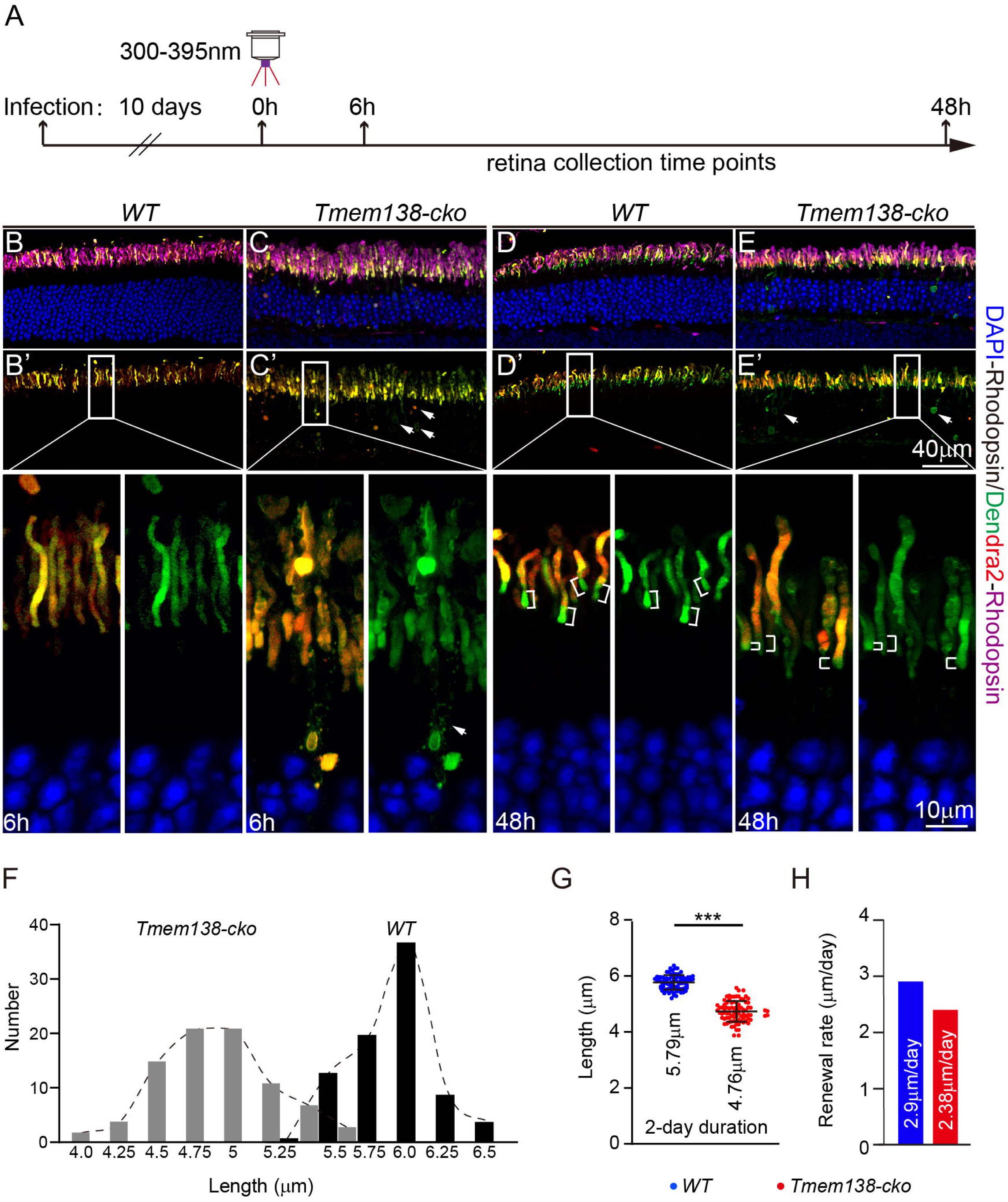
Photoreceptor OS renewal revealed by Rhodopsin/Dendra2 photoconversion. (**A**) Experimental scheme for detection of OS renewal at different time points after viral infection and UV radiation. Photoconversion was carried out on D10 post viral infection using a 300-395nm bandpass filter (Zeiss filter set 49) at 120 lux; 31 μw/cm^2^ illumination intensity and was observed 6 and 48 hours later. (**B-E**) Merged images from endogenous rhodopsin (magenta) and unconverted Rhodopsin/Dendra2 (green) and photoconverted (red) fluorescence of wild type (**B**, **D**) and *Tmem138*-*cko* mutant photoreceptors (**C**, **E**). (**B’**-**E’**) Merged unconverted (green) and photoconverted Rhodopsin/Dendra2 (red) fluorescence of the same set of retinal sections from (**B-E**). Magnified inset micrographs are displayed below as merged (left) and a green channel alone from (**B’**-**E’**). Brackets indicate the newly synthesized unconverted Rhodopsin/Dendra2 domains. (**F**) Distribution of newly grown lengths at the base of the wild type (dark gray boxes) and *Tmem138*-*cko* (light gray boxes) OS measured 48 hours after photoconversion with a 0.25μm bin width. (**G**) Student’s t-test was used to evaluate the significance of differences in average grown lengths between wild type and mutant OSs. ***, p<0.001. (**H**) Calculated growing rates based on unconverted pure fluorescent domains.

**Table 3.** Distribution of photoreceptors with differential Rhodopsin/Dendra2 domain lengths 48 hours after photoconversion.

| Length (μm) \ No. of PR | *WT* | *Tmem138-cko* |
| --- | --- | --- |
| 3.75-4 | 0 | 2 |
| 4-4.25 | 0 | 4 |
| 4.25-4.5 | 0 | 15 |
| 4.5-4.75 | 0 | 21 |
| 4.75-5 | 0 | 21 |
| 5-5.25 | 1 | 11 |
| 5.25-5.5 | 13 | 7 |
| 5.5-5.75 | 20 | 3 |
| 5.75-6 | 37 | 0 |
| 6-6.25 | 9 | 0 |
| 6.25-6.5 | 4 | 0 |
| Mean (μm) | 5.797 | 4.76 |
| Sd | 0.2565 | 0.3698 |

The left column number ranges are binned at 0.25μm intervals. No. PR, number of photoreceptors ; Sd: Standard deviation

### Evaluation of OS renewal with Peripherin2/Dendra2

To test a second membrane protein as a possible reporter for tracking OS renewal, we applied the same strategy to peripherin 2, a protein enriched in the OS disc rim, using a Peripherin2/Dendra2 fusion construct. A C-terminal tagged peripherin 2 using Dendra2 has been shown to have normal photoreceptor IS/OS localization in Xenopus photoreceptors^45^. While we observed largely normal localization of Peripherin2/Dendra2 in OS, there was a small amount of Peripherin2/Dendra2 mislocalized in the outer nuclear layer (ONL) at D4 and D6 post infection in both wild type and *Tmem138* mutant photoreceptors (Fig. 7A-D). OS localized Peripherin2/Dendra2 appeared to have an uneven banded pattern (Fig. 7A-D’’), similar to the observations by Hsu et al (2015)^46^. ONL localized protein was cleared in both wild type and mutant photoreceptors by10 days’ post-infection, and most photoreceptor OSs were filled with Peripherin2/Dendra2 along their entire length with a banded distribution pattern (Fig. 7E-F”).

**Figure 7.**
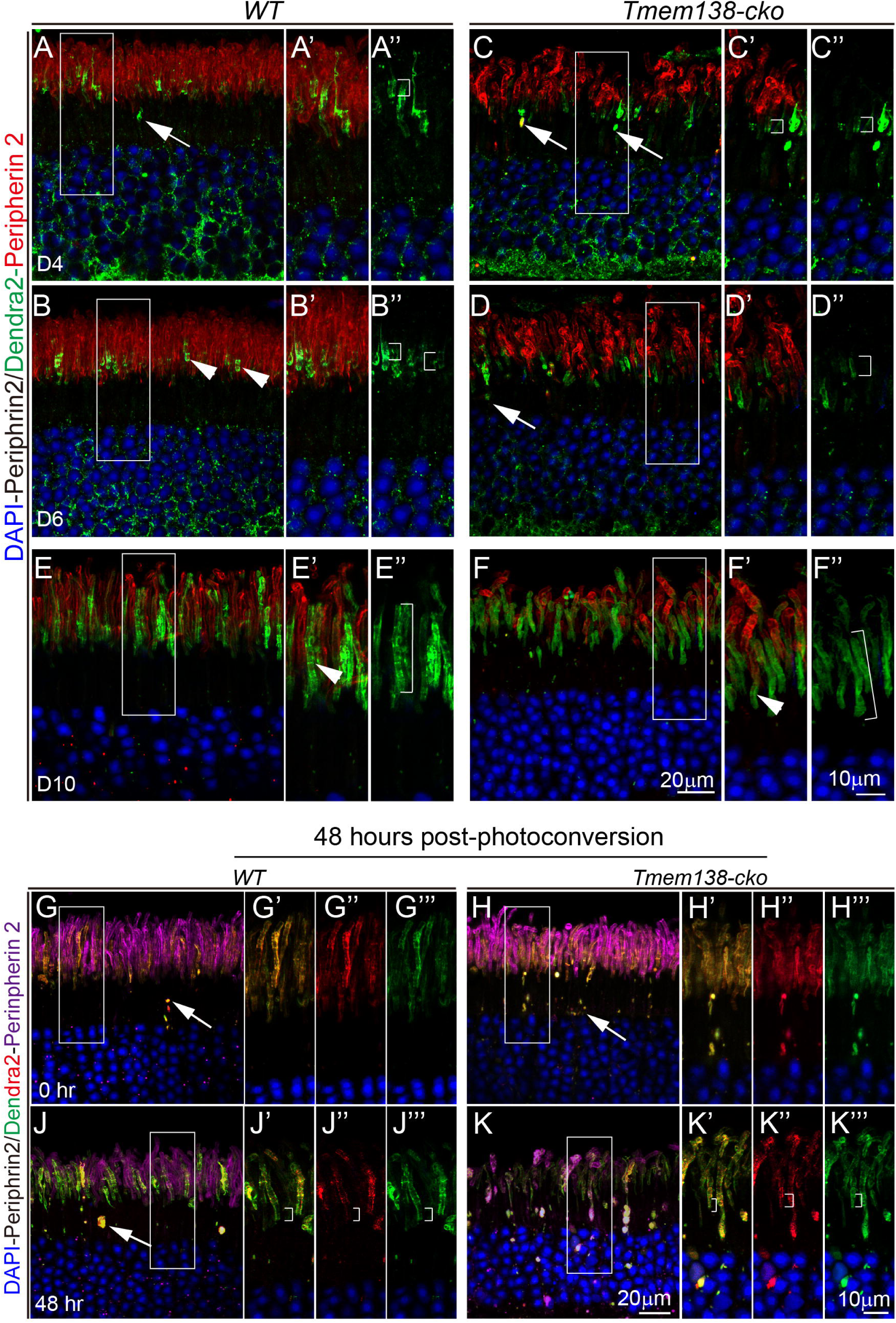
An evaluation of OS renewal with Peripherin2/Dendra2. For all panels, arrows point to the inner segment mislocalized Peripherin2/Dendra2, whereas arrowheads indicate the banded distribution pattern Peripherin2/Dendra2 within the OS. Brackets denote the approximate lengths of presumable renewed discs. Higher-magnification views of boxed regions are shown to the right of each corresponding panel. (**A-A**”) and (**B-B**”), Images taken from wild type retinal sections at D4 and D6 post infection, respectively. Endogenous peripherin 2 was labeled red with a specific antibody. (**C-C**”) and (**D-D**”), *Tmem138-cko* retinal sections at D4 and D6 post-infection, respectively, stained and imaged under the same conditions as wild type controls. (**E-E’**) and (**F-F**”), Wild type and *Tmem138-cko* retinal sections at D10 post infection. Note that the unconverted Peripherin2/Dendra2 was distributed along the full-length of the OSs. (**G-H’”**), Retinal sections prepared immediately after photoconversion, showing successful photoconversion with overlapping green (unconverted) and red (converted) Dendra2 signals. Endogenous peripherin 2 was labeled in magenta using immunostaining (see Materials and Methods). (**J-K’”**) Retinal sections taken 48 hours post-photoconversion. Note the lack of a distinct basal green-only zone at the OS base, the intermingled red/green fluorescence, and increased inner segment mislocalization. The green signal also exhibits a banded distribution pattern. To prevent unintended secondary photoconversion during imaging, the DAPI channel was acquired last in all image sets.

We next performed photoconversion in both wild type and mutant photoreceptors after Peripherin2/Dendra2 had occupied the entire OS. Successful photoconversion was confirmed on retinal sections prepared immediately after UV illumination, as indicated by abundant red Peripherin2/Dendra2 in the OS (Fig. 7G-H”’). However, unlike the distinct, basal-restricted green-only zones observed with Rhodopsin/Dendra2 at the same time point, Peripherin2/Dendra2 signals were not restricted to the basal zones. Instead, green and red fluorescence intermingled throughout the length of the OS (Fig. 7J–K□). Peripherin2/Dendra2 also showed increased mislocalization to the inner segments in both wild type and *Tmem138* mutant photoreceptors two days following UV exposure (Fig. 7J–K″), and ONL mislocalization was more evident in the mutant photoreceptors (Fig. 7K, K’-K’’’). Collectively, our data indicates that under the current experimental settings, C-terminus tagged peripherin 2 by Dendra2 is not a suitable reporter for OS renewal.

## Discussion

Renewal of the OS is a distinct aspect of photoreceptor cell biology, a process that requires directional trafficking of large amount of proteins and lipids to support new disc formation. Numerous disease-causing mutations in the retina have already been implicated in OS renewal, and many more likely remain to be discovered. Methods that allow direct visualization of OS renewal are critical for advancing the basic and translational studies in this area. While photoreceptor OS renewal has been extensively studied in amphibians using photoconvertible fluorescence tags^15, 34–36, 45^, direct visualization of rhodopsin OS transport and renewal in mice poses unique challenges. Several factors contribute to these difficulties: 1) Mouse photoreceptor OS is significantly smaller in diameter—approximately one-quarter the size of that in frogs^37, 42, 47, 48^—and photoreceptors are densely packed within the outer retina. This dense arrangement complicates the visualization of individual OS structures; 2) Distinguishing newly synthesized disc membrane proteins from preexisting ones is critical for monitoring OS transport over time. However, tools offering such temporal resolution have been lacking in mice; 3) Unlike amphibians, introducing an exogenous reporter gene into the mouse genome is a lengthy process. Crossing reporter lines into mutant backgrounds with defective OS transport further prolong the timeline.

To address these challenges, we utilized AAV-mediated delivery of photoconvertible Dendra2-tagged OS proteins into the somatic retina. This approach circumvents the time-consuming process of germline transgenics and provides a flexible and efficient platform for studying photoreceptor renewal and disc protein transport in both physiological and diseased states. AAV-delivered Dendra2-tagged rhodopsin serves as a fluorescent marker that can approximate the OS renewal timeline, particularly when its expression occurs within the first OS renewal cycle. However, calculating renewal rates by measuring the fluorescent domain lengths at two time points is subject to the inherent variability in expression onset and levels from viral transduction. Recombinant protein expression from AAV vectors exhibit a significant and variable delay relative to plasmid transfection, thought to be due to the time required for second strand DNA synthesis before the vector DNA becomes transcriptionally competent. Nevertheless, this approach can roughly distinguish the OS renewal rates between wild type and *Tmem138* mutant photoreceptors.

Once the Dendra2-tagged rhodopsin expression exceeds the duration of a full renewal cycle, photoconversion provides a far more reliable means of resetting the renewal timeline to “time zero.” By converting all preexisting rhodopsin fluorescence to red, any subsequently observed pure green fluorescence can, in theory, be unequivocally attributed to newly synthesized rhodopsin in the outer segment. This approach eliminates the variability brought on by expression onset from AAV vectors and thus enables precise tracking of OS renewal from the very onset.

Technical concerns include whether photoconvertible proteins faithfully recapitulate the behavior of their unmodified, endogenous proteins and whether UV irradiation might damage the retina, potentially impairing OS renewal. Our results indicate that Dendra2-tagged rhodopsin exhibits normal OS localization, and the measured OS renewal rate closely aligns with those previously reported, i.e., approximately 8–10% of the OS length per day^37^. Furthermore, a reduced OS renewal rate was observed in *Tmem138* mutant photoreceptors, which have compromised OS morphogenesis and protein trafficking^24^. This method, in principle, is suitable to uncover OS renewal defect in mammalian models. Other limitations, such as variability in AAV expression timing and delivery efficiency, may also warrant consideration. Regarding the former, once AAV vectors are fully expressed, photoconversion effectively eliminates the impact of timing variability. The latter is less concerning, as AAV has been extensively characterized as a robust and efficient expression vector in the retina. Moreover, by adjusting the viral titer or modifying the injection method, either localized or widespread retinal expression can be readily achieved.

Several methods have been developed over the years to track photoreceptor outer segment (OS) renewal in mice. The earliest work by Young employed radiolabeled amino acids, which provided limited spatial resolution^1^. More recently, a water-soluble and membrane-impermeable fluorescent dye, CF-568-hydrazide was used to monitor OS renewal through labeling of closed discs via intravitreal injection, yielding promising results^49^. While this approach has the advantages of reducing retinal phototoxicity and ease of application, it might be difficult to precisely define the initiation time point of renewal due to uncertain diffusion kinetics of the dye from the vitreous to the OS. Consequently, the method relies on labeling at two separate time points of injections within a single eye^49^. Moreover, the spatial resolution may be compromised by the dye’s water solubility and membrane permeability. In contrast, using an intrinsic OS membrane protein such as rhodopsin as a renewal marker, combined with photoconversion to “reset” the renewal clock, enables real-time evaluation of OS dynamics and thus expands the available toolkit for studying photoreceptor renewal. A direct comparison between our methodology and other related approaches has been summarized in Table 4.

Unlike rhodopsin, Peripherin2/Dendra2 failed to track OS renewal, despite appearing—at least partially—correctly localized (Fig. 7G, H). However, ONL mislocalization of Peripherin2/Dendra2 was observed, particularly at early ages following AAV infection. This may reflect the photoreceptor’s gradual adaptation to trafficking overexpressed exogenous proteins. Additionally, inner segment mislocalization of Peripherin2/Dendra2 may result from its intrinsic biochemical properties since this does not happen in tagged rhodopsin construct. These factors combined might partly contribute to the absence of clearly separated red (converted) and green (nascent) fluorescence domains in the OS, which warrant further investigation in future studies.

Although speculative, intrinsic features of peripherin 2 trafficking as suggested by earlier studies^45, 46, 50^ may also limit its suitability for tracking the OS renewal. Indeed, alternating stacks of rhodopsin- and peripherin 2-rich discs within OS have been reported^46^. Our previous study also showed that peripherin 2 and rhodopsin were not completely colocalized at least on postnatal day 8 (Guo et al., 2022^24^ supplementary data) suggesting asynchronous integration dynamics between the two proteins. Many other disc proteins may also exhibit distinct membrane integration or transport behaviors precluding them from being an optimal tracker for OS renewal.

Our study is the first to demonstrate successful application of Dendra 2 fusion protein in tracking OS renewal in mice through AAV-mediated gene delivery. This method was validated by revealing distinct OS renewal rates between wild type and *Tmem138* mutant photoreceptors, a ciliopathy model. The spatiotemporal localization of Dendra2-tagged rhodopsin along with its photoconverted form allows for “real-time” and direct estimation of the OS renewal alterations in diverse photoreceptor disease models. As OS renewal defects and rhodopsin mislocalization in many germline mutants may be secondary to severe photoreceptor degeneration, this method is particularly useful in late onset retinal degeneration models where OS structure and rhodopsin localization remain relatively preserved. Alternatively, when combined with temporally controlled genetic deletion, such as using Cre-ER as driver, this approach can be leveraged to investigate acute changes in OS renewal or rhodopsin transport. Lastly, although not yet feasible, with future advancements in high-resolution live imaging technologies, our method could potentially be applied to clinical diagnosis and to monitoring therapeutic outcomes in retinal diseases.

## MATERIALS AND METHODS

### Animals

The Tmem138 null (*Tmem138^b/b^*) and conditional(*Tmem138^b/c^; Rho-Cre*) mutant mice were generated as previously described^24^, and are referred to here as *Tmem138^−/-^* and *Tmem138-cko*, respectively, for simplicity. All animals were housed in a specific pathogen-free facility with 12:12-hour light/dark cycle. All experimental procedures involving mice were approved by the Zhongshan Ophthalmic Center Animal Care and Use Committee (ACUC).

### Dendra2-tagged constructs, lentiviral preparation, *in vitro* photoconversion, confocal microscopy, and measurement of fluorescence intensity

The full-length mouse *Rhodopsin* cDNA was fused to *Dendra2* at C-terminus and cloned into the pLVX vector with a CMV promotor. The resulting *Rhodopsin/Dendra2* construct was packaged into lentiviral particles using a third-generation lentiviral system consisting of the packaging plasmids pMD2.G, pMDLg/pRRE, and pRSV-Rev, co-transfected into HEK 293T cells. Viral supernatants were collected, filtered, and concentrated by ultracentrifugation at 100,000 × g for 2 hours at 4°C. Pelleted viral particles were resuspended in 500 μl of DMEM, aliquoted, and stored at −80°C until use.

Fifty microliters of packaged Rhodopsin/Dendra2 lentivirus were used to infect 293T cells cultured in glass-bottom confocal dishes (801002, NEST, USA) in the presence of polybrene (H9268, Sigma, USA) at a final concentration of 4 μg/ml. After a 48-hour infection, live 293T cells expressing Rhodopsin/Dendra2 were subjected to photoconversion using ultraviolet (UV) light banded for 300-395 nm (Zeiss DAPI filter set 49: G365; FT395; BP445/50) (Fig. 1C). Illumination was delivered through a 40× objective lens, with an intensity measured at 164 lux (39□μW/cm²) using an X-Cite 120Q mercury light source (EXCELITAS, USA).

For confocal microscopy, Dendra2 fluorescence was captured both before and after photoconversion using defined laser and detector settings. Green (unconverted) Dendra2 was excited with a 488nm gallium arsenide laser and detected within 493-550nm emission range. Red (photoconverted) Dendra2 was excited using a 561□nm laser line, with emission collected in the 568–640□nm range (Fig. 1C). The same laser and detector settings were also applied to immunofluorescence imaging of tissue sections.

Fluorescence intensities were measured from four images using NIH ImageJ software (arbitrary units). Green fluorescence signals were normalized to the average maximum intensity of unconverted Rhodopsin/Dendra2. To standardize graphing scales, red fluorescence intensities were normalized to 80% of the maximum green intensity, reflecting the ∼20% residual unconverted green fluorescence observed under the photoconversion conditions used.

### AAV production

Full-length mouse *Rhodopsin* and *Peripherin 2* cDNAs, each fused with *Dendra2* at the C-terminus, were cloned into a AAV2/8 vector driven by a human rhodopsin kinase (*hRK*) promoter^43^. The resulting constructs were co-transfected into HEK 293T cells along with plasmids encoding the AAV rep and cap genes (pAAV-RC8) and the adenoviral helper genes (pAdHelper), using Lipofectamine 2000 (11668027, Thermo Fisher Scientific, USA). Forty-eight hours post-transfection, the culture supernatant was harvested and AAV particles were purified using an iodixanol gradient ultracentrifugation protocol (https://www.addgene.org/protocols/aav-purification-iodixanol-gradient-ultracentrifugation/). Briefly, the supernatant was cleared by low-speed centrifugation to remove cellular debris, followed by ultracentrifugation through a 10% to 40% iodixanol (D1556, Sigma, USA) gradient at 150,000 × g for 6 hours. AAV-containing fractions were then collected and subjected to buffer exchange from iodixanol to PBS using Amicon Ultra-15 Centrifugal Filter Units (UFC905008, Millipore, USA). Viral titers were quantified using quantitative PCR with ITR (inverted terminal repeat) primers.

### Subretinal injections

Four-week-old mice were anesthetized via intraperitoneal injection of ketamine (80 mg/kg) and xylazine (4 mg/kg), followed by topical application of 1% tropicamide and 2.5% phenylephrine to induce pupil dilation. Mice were placed on a heating pad and positioned under an operating microscope (SZX7, Olympus, Japan). A drop of phosphate-buffered saline (PBS) was applied to the cornea to create a convex lens effect, enhancing fundus visualization.

A guide incision at the limbus was made using a 26-gauge needle inserted at a 45° angle toward the optic nerve. A microinjection syringe fitted with a 33-gauge blunt tip needle (7762-06, Hamilton, USA) was then used to deliver 0.5-1 µl of AAV containing approximate 5×10^8^ viral particles into the subretinal space over ∼ 5 seconds. Successful delivery was confirmed by the appearance of a retinal bleb under microscopy. After injection, mice were allowed to recover on the heating pad before being returned to their cages.

### In vivo photoconversion and imaging

AAV-infected mice were maintained to post-injection time points of day 4, 6 and 10. Mice were anesthetized and pupil-dilated as previously described. Animals were placed under an epifluorescence microscope equipped with 4x objective lens (Imager.Z2, Zeiss, Germany). A drop of PBS was applied to each eye to create a convex lens effect facilitating retinal visualization. The microscope was adjusted to focus on the retina guided by the fundus blood vessels. Ultraviolet (UV) light for photoconversion was delivered using a mercury arc lamp (X-Cite 120Q, EXCELITAS, USA) with a five-step iris diaphragm. A Zeiss DAPI filter set 49 (excitation: G365; beam splitter: FT395; emission: BP445/50) was used to isolate UV wavelengths ranging from 300–395□nm, with a peak emission at 365□nm (Fig.□2B).

UV irradiance and visible illuminance were measured separately by placing either an Ultraviolet Irradiance Meter (UV-A, HANDY, China) or an Illuminance Meter (PP710, SanLiang, China) 1 centimeter below a 4× objective — a distance optimized for photoconversion. Rhodopsin/Dendra2 photoconversion was assessed under four UV intensities—40 lux/5 μW/cm², 66 lux/18 μW/cm², 120 lux/31 μW/cm², 390 lux/96 μW/cm² —across three exposure durations: 5, 10, and 15 minutes.

For image collection, unconverted green and photoconverted red Dendra2 fluorescence were detected using Zeiss filter set 38 for GFP (BP470/40; FT495; BP 525/50) and Zeiss filter set 20 for RFP (BP 546/12; FT 560, BP 575-640), respectively (Fig. 2F, G).

### Retinal dissection, sectioning, immunostaining, and imaging

Following rapid euthanasia, mouse eyeballs were immediately enucleated and submerged in cold PBS for dissection. The cornea and sclera were carefully removed, followed by lens removal. Isolated retinas were fixed in 4% paraformaldehyde (PFA) containing 0.25% glutaraldehyde for 30 minutes at room temperature. Retinas were thoroughly washed in PBS to eliminate residual fixatives and embedded in 4-5% agarose for vibratome sectioning. Retinal sections of 100 µm thickness were prepared using a Leica Vibratome (VT1000S, Leica, Germany).

For immunostaining, sections were blocked with PBST (PBS + 0.5% Triton X-100) with 5% donkey serum at room temperature for 30 minutes. Sections were then incubated with primary antibodies against rhodopsin (1D4, 1:500, a gift from Dr. Tiansen Li at NEI) or peripherin 2/Rds (ab122057, Abcam, USA) at 4°C for at least 24 hours. After three extensive washes with PBST, sections were incubated with Alexa Fluor 568 or Alexa Fluor 647-conjugated secondary antibodies in blocking buffer at 4°C for additional 24 hours. Retinal slices were washed in PBST and rinsed with PBS before mounting on glass slides for imaging.

Imaging was performed using a Zeiss LSM880 confocal microscope equipped with an Airyscan detector. The following laser lines and detection settings were used for each fluorophore:

Alexa Fluor 647: Excitation with 633□nm laser; main beam splitter (MBS): 488/561/633; detection filter: BP 570–620□nm + LP 645□nm; Dendra2 (red): Excitation with 561□nm laser; MBS: 458/561; detection filter: BP 495–535□nm + LP 555□nm; secondary beam splitter (SBS): SP 615; Dendra2 (green): Excitation with 488□nm laser; MBS: 488; detection filters: BP 420–480□nm + BP 495–550□nm; DAPI: Excitation with 405□nm laser; MBS: 405; detection filters: BP 420–480□nm + BP 495–550□nm

### Quantification and statistics

For HEK293T cells, Dendra2 fluorescence intensities were quantified from images captured at four different locations using a 40×/1.4 oil objective. Total fluorescence intensities were calculated using ImageJ and normalized to measured areas (per 100 μm²). Unconverted green fluorescence intensities at different time durations were normalized to the initial intensity, while the converted red fluorescence intensities were normalized to that of the longest time of exposure (120s). Data was presented as a percentage.

For measuring the lengths of rhodopsin or peripherin 2/Rds trafficking, images were acquired using a 100×/1.4 oil objective equipped on Zeiss LSM880 with Airyscan. Length measurement was conducted using 100-150 rod outer segments from each retina section (n=3 animals) and analyzed by NIH ImageJ software.

All statistical analyses were conducted using GraphPad Prism 7.0 software and presented as mean ± standard deviation (SD). Statistical significance was determined using the Student’s t-test, with results denoted as follows: “*”, p < 0.05; “**”, p< 0.01, “***”, p < 0.001.

