## Supplementary Figures and Tables for "Visualization of photoreceptor outer segment renewal using AAV-delivered Dendra2-tagged rhodopsin"

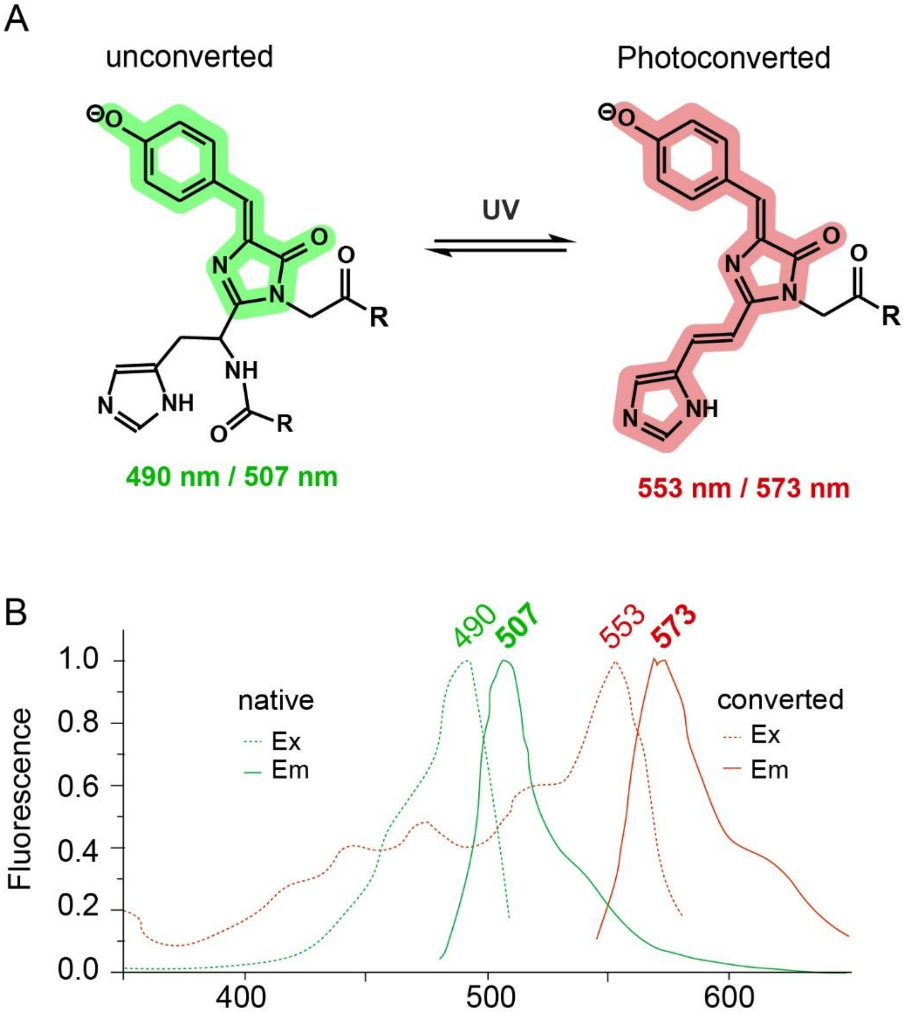

**Supplementary Figure 1**. **Dendra2 photoconversion and associated excitation/emission spectra.** (**A**), Schematic illustration of Dendra2 chromophore transformations under UV illumination (Remade from Pakhomov et al., 2017., reference 31) (**B**), Excitation (dashed lines) and emission (solid lines) spectra of Dendra2 before (green) and after (red) photoconversion (remade from Chudakov et al., 2007, reference 40).

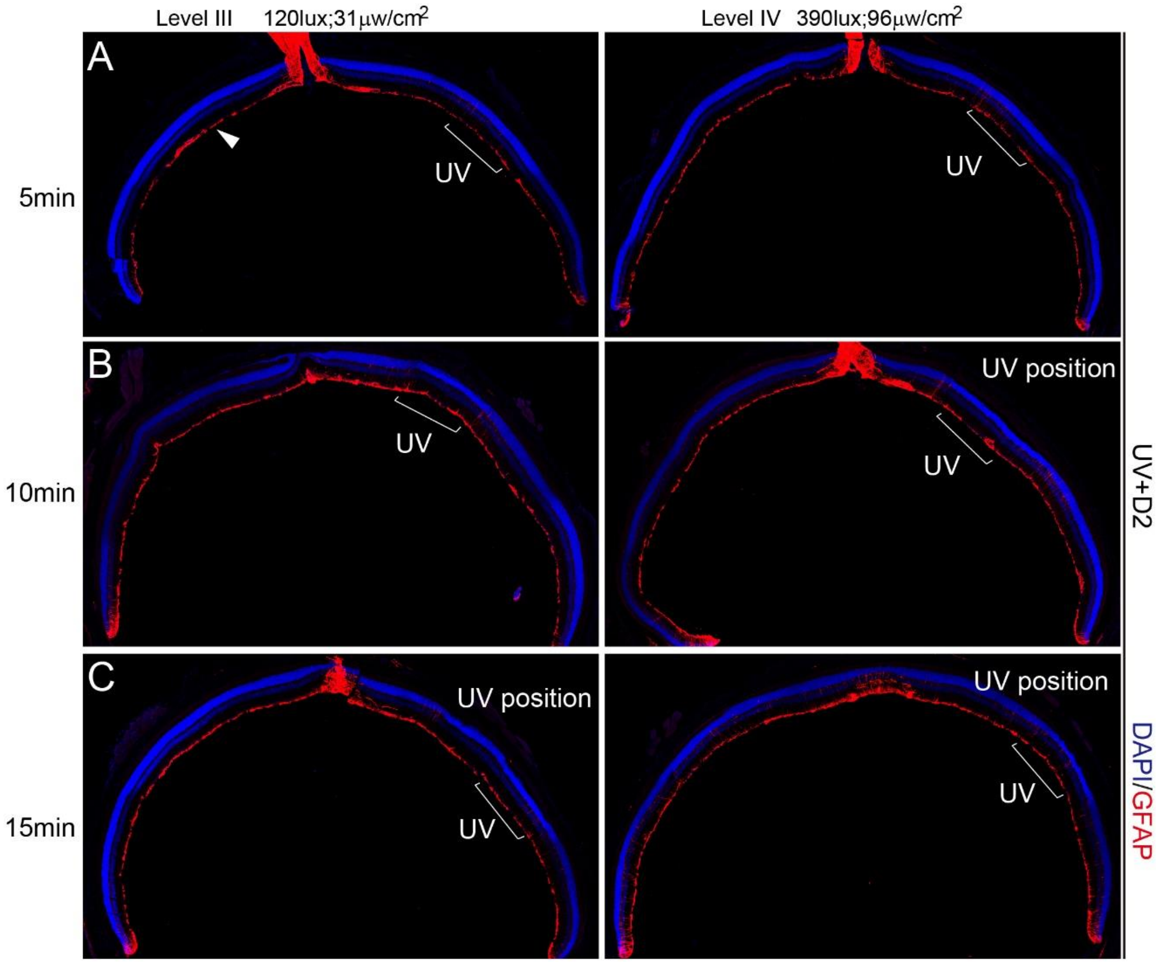

**Supplementary Figure 2**. **Evaluation of retinal light damage with different illumination intensities over time.** Retinal sections were stained for GFAP (red) and DAPI (blue) two days after UV illumination (UV+ D2). (**A**) Level III and IV illumination for 5min only minimally altered GFAP activation and retinal structure. (**B, C**) Level III and IV illumination for 10 and 15 minutes, respectively, resulted in loss of the outer nuclear layer and widespread GFAP activation.

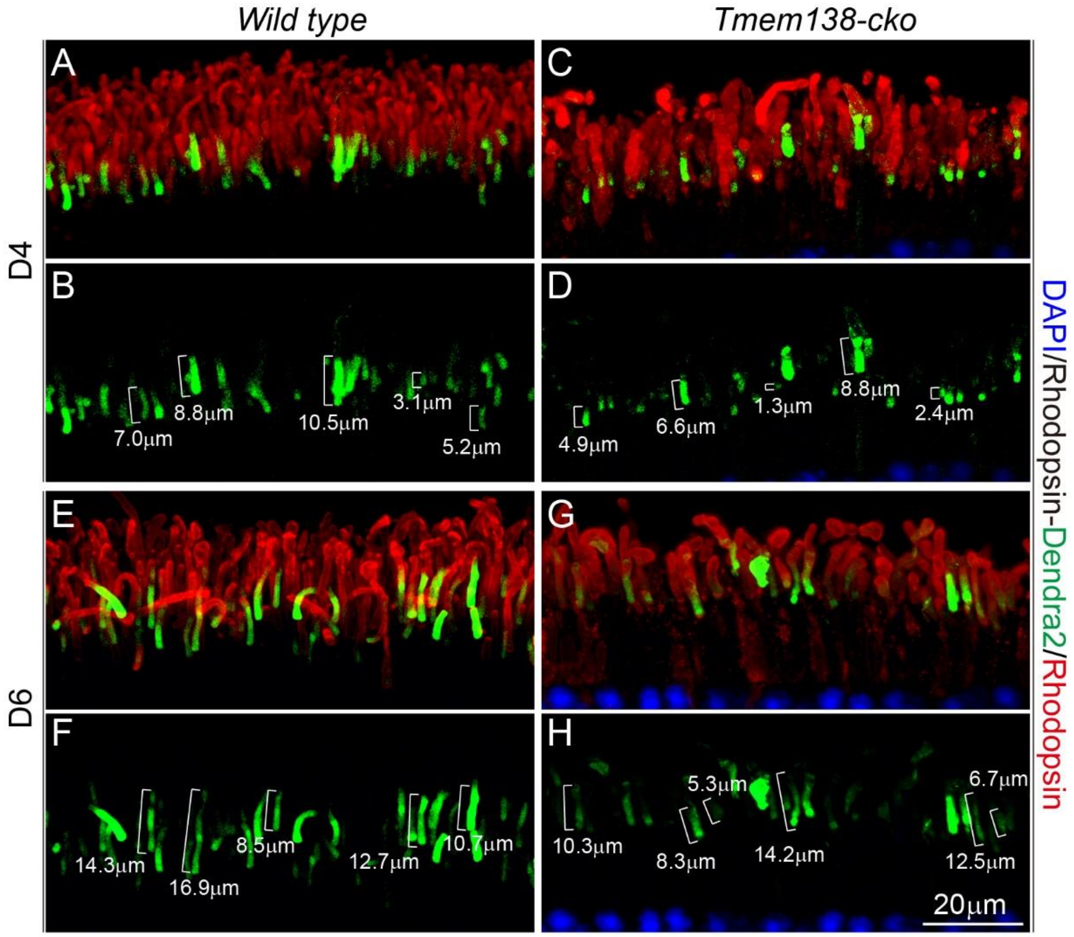

**Supplementary Figure 3**. **Rhodopsin/Dendra2 expression and localization during OS renewal.** Endogenous Rhodopsin was labeled with ID4 antibody (red) and AAV-introduced Rhodopsin/Dendra2 was in green. Brackets in (**B**, **D, F, H**) refer to measured Rhodopsin/Dendra2 domain lengths. (**A, B**), Wild type retinal sections at D4 post infection. (**A**)**,** Merged image; (**B**), Green channel alone. (**C, D**), *Tmem138-cko* retinal sections at D4 post infection. Same panel arrangement as in (**A**, **B**). (**E, F**), Wild type retinal sections at D6 post infection with same panel arrangement as in (**A**, **B**). (**G, H**), *Tmem138-cko* retinal sections at D6 with same panel arrangement as in (**E**, **F**). Note shortened Rhodopsin domains of the *Tmem138-cko* mutant photoreceptors at either D4 or D6.

**Supplementary Table 1. Measured Rhodopsin/Dendra2 domain length of each photoreceptor for Table 2 and Table 3**

| Length (μm) / #PR | Unconverted photoreceptors —Table 2 | | | | 48h after conversion —Table 3 | |
|---|---|---|---|---|---|---|
| | WT_D4 | WT_D6 | Tmem138 -cko_D4 | Tmem138 -cko_D6 | WT | Tmem138 -cko |
| 1 | 2.4 | 8.2 | 0.9 | 5.8 | 5.41 | 4.71 |
| 2 | 2.6 | 8.3 | 0.8 | 5.9 | 5.23 | 4.22 |
| 3 | 3.2 | 8.5 | 1.6 | 5.4 | 5.97 | 4.60 |
| 4 | 3.3 | 8.3 | 1.4 | 5.3 | 5.65 | 4.50 |
| 5 | 3.5 | 9.1 | 1.2 | 5.8 | 5.56 | 4.82 |
| 6 | 4.4 | 9.3 | 2.1 | 6.9 | 5.62 | 4.60 |
| 7 | 4.3 | 9 | 2.3 | 6.7 | 5.41 | 4.46 |
| 8 | 4.6 | 9.7 | 2.6 | 6.5 | 5.34 | 4.89 |
| 9 | 4.2 | 9.8 | 2.4 | 6.6 | 6.02 | 4.41 |
| 10 | 4.9 | 9 | 2.9 | 6.8 | 6.17 | 5.09 |
| 11 | 4.3 | 9.2 | 2.8 | 6.2 | 5.50 | 5.08 |
| 12 | 4.2 | 9.7 | 2.4 | 6.3 | 5.94 | 4.56 |
| 13 | 4.5 | 10.7 | 2.1 | 6.9 | 5.74 | 4.89 |
| 14 | 4.8 | 10.9 | 2.2 | 7.3 | 5.53 | 5.07 |
| 15 | 5.1 | 10.4 | 2.3 | 7.4 | 5.47 | 4.29 |
| 16 | 5.2 | 10.3 | 3.6 | 7.5 | 5.64 | 4.69 |
| 17 | 5 | 10.1 | 3.2 | 7.2 | 5.92 | 4.62 |
| 18 | 5.5 | 10.7 | 3.6 | 7.7 | 5.66 | 4.59 |
| 19 | 5.9 | 10.6 | 3.7 | 7.6 | 5.53 | 4.85 |
| 20 | 5.1 | 10.4 | 3.9 | 7.4 | 5.87 | 4.87 |
| 21 | 5.3 | 10.1 | 3.6 | 7.1 | 5.83 | 4.69 |
| 22 | 5.6 | 10.4 | 3.1 | 7.9 | 5.77 | 5.18 |
| 23 | 5.7 | 10.7 | 3.4 | 7.8 | 5.99 | 5.19 |
| 24 | 5.9 | 10.1 | 3.5 | 7.6 | 6.08 | 5.17 |
| 25 | 5.3 | 11.3 | 3.3 | 7.7 | 5.44 | 4.85 |
| 26 | 5.1 | 11.4 | 3.7 | 7.2 | 5.91 | 4.59 |
| 27 | 5.8 | 11.5 | 3.4 | 7.4 | 5.59 | 4.87 |
| 28 | 5.2 | 11.2 | 3.8 | 7.9 | 5.96 | 4.89 |
| 29 | 5.6 | 11.7 | 3 | 7.2 | 5.56 | 5.10 |
| 30 | 5.5 | 11.6 | 3.4 | 8.2 | 6.17 | 4.59 |
| 31 | 6.2 | 11.4 | 3.6 | 8.3 | 5.31 | 4.41 |
| 32 | 6.3 | 11.1 | 3.7 | 8.5 | 6.09 | 4.53 |
| 33 | 6.4 | 11.9 | 3.3 | 8.3 | 5.95 | 5.29 |
| 34 | 6.1 | 11.8 | 3.1 | 8.8 | 5.84 | 5.32 |

| | | | | | | |
|---|---|---|---|---|---|---|
| 35 | 6.6 | 11.6 | 4.8 | 8.1 | 5.49 | 4.89 |
| 36 | 6.5 | 11.7 | 4.9 | 8.3 | 5.77 | 5.51 |
| 37 | 6.3 | 11.2 | 4.4 | 8.6 | 5.56 | 5.07 |
| 38 | 6 | 11.3 | 4.3 | 8.4 | 6.17 | 4.46 |
| 39 | 6.9 | 11.4 | 4.8 | 8.9 | 5.59 | 4.41 |
| 40 | 6.7 | 11.9 | 4.4 | 8.8 | 5.87 | 4.53 |
| 41 | 6.5 | 11.2 | 4.9 | 8.4 | 5.47 | 5.22 |
| 42 | 6.6 | 11.1 | 4.2 | 8.1 | 5.72 | 5.29 |
| 43 | 6.1 | 11.5 | 4.9 | 8.2 | 5.98 | 5.51 |
| 44 | 6.2 | 11.9 | 4.8 | 8.3 | 6.30 | 4.89 |
| 45 | 6.3 | 11.2 | 4.1 | 8.9 | 5.95 | 5.07 |
| 46 | 6.8 | 11.6 | 4.3 | 8.4 | 5.43 | 4.21 |
| 47 | 6.1 | 12.7 | 4.6 | 8.3 | 5.49 | 5.21 |
| 48 | 6 | 12.8 | 4.1 | 8.4 | 5.91 | 4.53 |
| 49 | 6.4 | 12.3 | 4.9 | 8.8 | 5.69 | 4.37 |
| 50 | 6.9 | 12.2 | 4.4 | 9.8 | 5.93 | 4.64 |
| 51 | 6.1 | 12.7 | 4.4 | 9.9 | 5.89 | 4.66 |
| 52 | 6.5 | 12.3 | 4.3 | 9.4 | 5.69 | 4.76 |
| 53 | 6.3 | 12.9 | 4.6 | 9.3 | 5.98 | 4.68 |
| 54 | 6.7 | 12.1 | 4.6 | 9.8 | 5.56 | 4.71 |
| 55 | 6.8 | 12.8 | 4.4 | 9.4 | 5.80 | 4.52 |
| 56 | 6.3 | 12.7 | 4.9 | 9.9 | 6.09 | 4.63 |
| 57 | 6.2 | 12 | 4.2 | 9.2 | 5.77 | 4.77 |
| 58 | 6.7 | 12.2 | 4.9 | 9.9 | 5.84 | 4.36 |
| 59 | 6.3 | 12.5 | 4.6 | 9.8 | 6.23 | 4.80 |
| 60 | 6.9 | 12 | 4.1 | 9.1 | 5.91 | 4.98 |
| 61 | 6.1 | 12.9 | 4.2 | 9.3 | 5.98 | 4.90 |
| 62 | 6.8 | 12.3 | 4.4 | 9.6 | 5.92 | 4.40 |
| 63 | 6.7 | 12.3 | 4.7 | 9.1 | 5.87 | 4.80 |
| 64 | 6 | 12.2 | 4.4 | 9.9 | 5.76 | 5.30 |
| 65 | 6.2 | 12.5 | 4.2 | 9.4 | 5.84 | 4.10 |
| 66 | 6.5 | 12.5 | 4.9 | 9.4 | 5.89 | 4.90 |
| 67 | 6 | 12.3 | 4.8 | 9.3 | 5.41 | 5.30 |
| 68 | 6.9 | 12.8 | 4.1 | 9.6 | 5.70 | 4.50 |
| 69 | 6.3 | 12.1 | 4.3 | 9.6 | 6.00 | 4.32 |
| 70 | 6.3 | 12.9 | 4.6 | 9.4 | 5.80 | 4.31 |
| 71 | 6.2 | 12.5 | 4.1 | 9.9 | 6.00 | 4.33 |
| 72 | 6.5 | 12 | 4.9 | 9.2 | 6.10 | 4.70 |
| 73 | 6.5 | 12 | 4.4 | 9.9 | 6.40 | 4.70 |
| 74 | 6.3 | 12.3 | 4.4 | 9.6 | 5.90 | 5.60 |
| 75 | 6.8 | 12.4 | 5.8 | 9.1 | 5.50 | 4.30 |
| 76 | 6.1 | 12.6 | 5.9 | 9.2 | 6.30 | 4.10 |

| | | | | | | |
|---|---|---|---|---|---|---|
| 77 | 6.9 | 12.9 | 5.4 | 9.4 | 5.90 | 5.00 |
| 78 | 6.5 | 12.1 | 5.3 | 9.7 | 5.80 | 4.80 |
| 79 | 6 | 13.2 | 5.8 | 9.4 | 6.30 | 5.30 |
| 80 | 7.1 | 13.9 | 5.4 | 9.2 | 6.00 | 3.90 |
| 81 | 7.3 | 13.8 | 5.9 | 9.9 | 5.60 | 3.90 |
| 82 | 7.4 | 13.1 | 5.2 | 9.8 | 5.61 | 5.30 |
| 83 | 7.5 | 13.3 | 5.9 | 9.1 | 5.60 | 5.00 |
| 84 | 7.2 | 13.6 | 5.8 | 9.3 | 6.00 | 4.90 |
| 85 | 7.7 | 13.1 | 5.1 | 10.1 | | |
| 86 | 7.6 | 13.7 | 5.3 | 10.3 | | |
| 87 | 7.4 | 13.4 | 5.6 | 10.6 | | |
| 88 | 7.1 | 13.4 | 5.1 | 10.1 | | |
| 89 | 7.2 | 13.3 | 5.9 | 10.9 | | |
| 90 | 7.8 | 13.6 | 5.4 | 10.4 | | |
| 91 | 7.6 | 13.6 | 5.4 | 10.4 | | |
| 92 | 7.7 | 13.4 | 5.3 | 10.3 | | |
| 93 | 7.2 | 13.9 | 5.6 | 10.6 | | |
| 94 | 7.3 | 13.2 | 5.6 | 10.6 | | |
| 95 | 7.4 | 13.9 | 5.4 | 10.4 | | |
| 96 | 7.9 | 13.6 | 5.9 | 10.9 | | |
| 97 | 7.2 | 13.1 | 5.2 | 10.2 | | |
| 98 | 7.1 | 13.2 | 5.9 | 10.9 | | |
| 99 | 7.5 | 13.3 | 5.6 | 10.6 | | |
| 100 | 7.1 | 13.1 | 5.1 | 10.1 | | |
| 101 | 7.2 | 13.6 | 5.2 | 10.2 | | |
| 102 | 7.6 | 13.9 | 5.4 | 10.3 | | |
| 103 | 7.4 | 13.2 | 5.7 | 10.1 | | |
| 104 | 7.8 | 13.4 | 5.4 | 10.6 | | |
| 105 | 7.9 | 13.7 | 5.2 | 10.7 | | |
| 106 | 7.4 | 13.8 | 5.9 | 10.2 | | |
| 107 | 7.3 | 13.4 | 5.8 | 10.4 | | |
| 108 | 7.8 | 13.4 | 5.1 | 10.7 | | |
| 109 | 7.4 | 13.2 | 5.3 | 10.8 | | |
| 110 | 7.9 | 13.9 | 5.6 | 10.9 | | |
| 111 | 7.2 | 13.3 | 5.1 | 10.4 | | |
| 112 | 7.9 | 13.7 | 5.9 | 10.1 | | |
| 113 | 7.8 | 13.6 | 5.4 | 10.6 | | |
| 114 | 7.1 | 13.3 | 5.4 | 10.2 | | |
| 115 | 7.3 | 13.2 | 5.3 | 10.7 | | |
| 116 | 7.6 | 13.4 | 5.6 | 10.3 | | |
| 117 | 7.1 | 13.9 | 5.6 | 10.1 | | |
| 118 | 7.9 | 13.3 | 5.4 | 10.6 | | |

| | | | | |
|---|---|---|---|---|
| 119 | 7.4 | 13.7 | 5.9 | 10.1 |
| 120 | 7.4 | 13.6 | 5.2 | 10.2 |
| 121 | 7.3 | 13.3 | 5.9 | 10.7 |
| 122 | 7.6 | 13.2 | 5.6 | 10.6 |
| 123 | 7.6 | 13.4 | 5.1 | 10.7 |
| 124 | 7.4 | 13.9 | 5.2 | 10.5 |
| 125 | 7.9 | 13.2 | 5.3 | 10.5 |
| 126 | 7.2 | 13.9 | 5.1 | 10.4 |
| 127 | 7.9 | 14.8 | 5.6 | 10.3 |
| 128 | 7.6 | 14.9 | 5.7 | 10.4 |
| 129 | 7.1 | 14.4 | 5.2 | 10.7 |
| 130 | 7.2 | 14.3 | 5.4 | 10.3 |
| 131 | 7.3 | 14.8 | 5.7 | 10.5 |
| 132 | 7.1 | 14.4 | 5.8 | 10.4 |
| 133 | 7.6 | 14.9 | 5.9 | 10.1 |
| 134 | 7.6 | 14.2 | 5.4 | 10 |
| 135 | 7.2 | 14.9 | 5.1 | 10.9 |
| 136 | 7.4 | 14.8 | 5.6 | 10 |
| 137 | 7.7 | 14.1 | 5.2 | 11.8 |
| 138 | 7.8 | 14.3 | 5.7 | 11.1 |
| 139 | 7.9 | 14.6 | 5.3 | 11.3 |
| 140 | 7.4 | 14.1 | 5.1 | 11.6 |
| 141 | 7.2 | 14.9 | 5.6 | 11.1 |
| 142 | 7 | 14.4 | 5.1 | 11.9 |
| 143 | 7.3 | 14.4 | 5.2 | 11.4 |
| 144 | 7.7 | 14.3 | 5.7 | 11.4 |
| 145 | 7.6 | 14.6 | 5.6 | 11.3 |
| 146 | 7.3 | 14.6 | 5.7 | 11.6 |
| 147 | 7.2 | 14.4 | 5.5 | 11.6 |
| 148 | 7.4 | 14.9 | 5.5 | 11.4 |
| 149 | 8.8 | 14.2 | 5.4 | 11.9 |
| 150 | 8 | 14.9 | 5.3 | 11.2 |
| 151 | 8.4 | 14.6 | 6.2 | 11.9 |
| 152 | 8.3 | 14.1 | 6.3 | 11.6 |
| 153 | 8.2 | 14.2 | 6.4 | 11.1 |
| 154 | 8.4 | 14.4 | 6.8 | 11.2 |
| 155 | 8.9 | 14.7 | 6.6 | 11.4 |
| 156 | 8.2 | 14.4 | 6.5 | 11.7 |
| 157 | 8.9 | 14.2 | 6.3 | 11.4 |
| 158 | 8.8 | 14.9 | 6 | 11.2 |
| 159 | 8.1 | 14.8 | 6.9 | 11.9 |
| 160 | 8.3 | 14.1 | 6.7 | 11.8 |

| | | | | |
|---|---|---|---|---|
| 161 | 8.6 | 14.3 | 6.5 | 11.1 |
| 162 | 8.1 | 14.6 | 6.6 | 11.3 |
| 163 | 8.9 | 14.1 | 6.8 | 11.6 |
| 164 | 8.4 | 14.9 | 6.2 | 11.1 |
| 165 | 8.4 | 14.4 | 6.3 | 11.9 |
| 166 | 8.3 | 14.4 | 6.9 | 11.4 |
| 167 | 8.8 | 14.3 | 6.8 | 11.4 |
| 168 | 8.7 | 14.6 | 6.1 | 11.3 |
| 169 | 8.4 | 14.6 | 6.3 | 11.6 |
| 170 | 8.9 | 14.4 | 6.6 | 11.6 |
| 171 | 8.2 | 14.9 | 6.1 | 11.4 |
| 172 | 8.9 | 14.2 | 6.9 | 11.9 |
| 173 | 8.6 | 14.9 | 6.4 | 11.2 |
| 174 | 8.1 | 14.6 | 6.4 | 11.9 |
| 175 | 8.2 | 14.1 | 6.3 | 11.6 |
| 176 | 8.4 | 14.2 | 6.6 | 11.1 |
| 177 | 8.7 | 14.3 | 6.6 | 11.2 |
| 178 | 8.4 | 14.1 | 6.4 | 11.3 |
| 179 | 9.7 | 14.6 | 6.9 | 11.1 |
| 180 | 9.6 | 14.7 | 6.2 | 11.6 |
| 181 | 9.4 | 14.2 | 6.9 | 11.7 |
| 182 | 9.4 | 14.4 | 6.6 | 11.2 |
| 183 | 9.3 | 14.7 | 6.1 | 11.6 |
| 184 | 9 | 14.8 | 6.2 | 11.2 |
| 185 | 9.4 | 14.9 | 6.3 | 11.6 |
| 186 | 9.1 | 14.4 | 6.1 | 11.7 |
| 187 | 9.9 | 15.3 | 7.3 | 11.9 |
| 188 | 9.2 | 15.4 | 7.4 | 11.6 |
| 189 | 9.1 | 15.5 | 7.5 | 11.1 |
| 190 | 9.7 | 15.2 | 7.2 | 11.4 |
| 191 | 9.4 | 15.7 | 7.7 | 11.5 |
| 192 | 9 | 15.6 | 7.6 | 11.3 |
| 193 | 9 | 15.4 | 7.4 | 11.7 |
| 194 | 9.1 | 15.1 | 7.1 | 11.4 |
| 195 | 10.6 | 15.6 | 7.9 | 11.8 |
| 196 | 10.7 | 15.8 | 7.8 | 11 |
| 197 | 10.1 | 15.6 | 7.6 | 11.4 |
| 198 | 10.6 | 15.7 | 7.7 | 11.6 |
| 199 | 10.3 | 15.2 | 7.2 | 11.7 |
| 200 | 10.4 | 15.3 | 8.2 | 11.3 |
| 201 | 10.7 | 15.4 | 8.3 | 11.1 |
| 202 | 10 | 15.9 | 8.5 | 11.2 |

| | | | | |
|---|---|---|---|---|
| 203 | | 15.2 | 8.3 | 11.8 |
| 204 | | 15.1 | 8.8 | 11.3 |
| 205 | | 15.5 | | 11 |
| 206 | | 15.7 | | 11.7 |
| 207 | | 15.2 | | 12.8 |
| 208 | | 15.6 | | 12.1 |
| 209 | | 15.4 | | 12.3 |
| 210 | | 15.8 | | 12.6 |
| 211 | | 15.9 | | 12.1 |
| 212 | | 15.4 | | 12.9 |
| 213 | | 15.3 | | 12.4 |
| 214 | | 15.8 | | 12.1 |
| 215 | | 15.4 | | 12.3 |
| 216 | | 15.8 | | 12.4 |
| 217 | | 16.2 | | 12.6 |
| 218 | | 16.3 | | 12.2 |
| 219 | | 16.4 | | 12.6 |
| 220 | | 16.8 | | 12.7 |
| 221 | | 16.6 | | 12.9 |
| 222 | | 16.5 | | 12.6 |
| 223 | | 16.3 | | 12.1 |
| 224 | | 16 | | 12.4 |
| 225 | | 16.9 | | 12.5 |
| 226 | | 16.7 | | 12.3 |
| 227 | | 16.5 | | 12.7 |
| 228 | | 16.6 | | 12.4 |
| 229 | | 16.8 | | 12.8 |
| 230 | | 16.2 | | 12 |
| 231 | | 16.3 | | 12.4 |
| 232 | | 16.8 | | 12.6 |
| 233 | | 17.3 | | 12.7 |
| 234 | | 17.8 | | 12.3 |
| 235 | | 17.4 | | 12.1 |
| 236 | | 17.3 | | 12.4 |
| 237 | | 17.6 | | 12.6 |
| 238 | | 17.9 | | 12.3 |
| 239 | | | | 13.9 |
| 240 | | | | 13.4 |
| 241 | | | | 13.4 |
| 242 | | | | 13.3 |
| 243 | | | | 13.6 |
| 244 | | | | 13.6 |

| | | | | |
|---|---|---|---|---|
| 245 | | | | 13.4 |
| 246 | | | | 13.9 |
| 247 | | | | 13.2 |
| 248 | | | | 13.9 |
| 249 | | | | 13.6 |
| 250 | | | | 13.1 |
| 251 | | | | 14.6 |
| 252 | | | | 14.5 |
| 253 | | | | 14 |
| 254 | | | | 14.1 |
| 255 | | | | 14.7 |
| 256 | | | | 14.3 |
